# Terminally differentiated CX3CR1^+^ natural killer cells drive coronary arteritis through an NFAT–T-bet axis

**DOI:** 10.64898/2026.11.20.767965

**Authors:** Masami Fujita, Hiroki Satooka, Shinsuke Hoshino, Shoma Matsumoto, Eric J. Verbeke, Kenta Kondo, Yasutoshi Agata, Masatsugu Ema, Yoshihiro Maruo, Takako Hirata

**Affiliations:** Department of Fundamental Biosciences, Shiga University of Medical Science, Otsu, Shiga 520-2192, Japan; Department of Pediatrics, Shiga University of Medical Science, Otsu, Shiga 520-2192, Japan; Department of Stem Cells and Human Disease Models, Research Center for Animal Life Science, Shiga University of Medical Science, Otsu, Shiga 520-2192, Japan; Premium Research Institute for Human Metaverse Medicine (WPI-PRIMe), Osaka University, Suita, Osaka 565-0871, Japan; Department of Biochemistry and Molecular Biology, Shiga University of Medical Science, Otsu, Shiga 520-2192, Japan

**Author notes:** These authors contributed equally (Masami Fujita, Hiroki Satooka).

## Abstract

Natural killer (NK) cells acquire potent effector function through terminal differentiation, yet the signals driving this program toward pathogenicity in vascular inflammation remain undefined. Using an FK565-induced mouse model of Kawasaki disease and single-cell transcriptomics from patients, we identify NK cells as major drivers of coronary arteritis through cytotoxicity and CCL5–CCR1-mediated recruitment of myeloid cells. This activity is concentrated in a terminally differentiated, T-bet-driven NK cell subset, the cardiac accumulation of which depends on CX3CR1, and a corresponding population is enriched in patient transcriptomic datasets. We further identify signaling through calcineurin and nuclear factor of activated T cells (NFAT) as an upstream driver of this program, with NFATc1 induced in NK cells. Calcineurin inhibition suppressed T-bet induction, reduced this subset and attenuated disease development. These findings define an NFAT–T-bet–CX3CR1 axis that couples terminal NK cell differentiation to coronary inflammation.

---

Kawasaki disease (KD) is an acute systemic vasculitis of childhood and the leading cause of acquired pediatric heart disease in developed nations^1,2^. Untreated, approximately 25% of patients develop coronary artery aneurysms (CAAs), which can cause myocardial infarction and sudden death later in life^3^. Early high-dose intravenous immunoglobulin (IVIG) has reduced CAA incidence to about 5%^4^, yet 10–20% of patients are IVIG-resistant and remain at high risk of irreversible cardiovascular damage^1^. Despite these clinical challenges, the cellular and molecular mechanisms underlying the preferential involvement of coronary arteries in KD remain poorly understood.

A major obstacle to elucidating these mechanisms is the profound scarcity of affected human cardiovascular tissue, making experimental animal models essential^5^. Three widely used models of KD-like vasculitis are induced by *Candida albicans* water-soluble fraction^6^, *Lactobacillus casei* cell wall extract^7^ or FK565, a nucleotide-binding oligomerization domain-containing protein 1 (NOD1) ligand^8^. Among these, systemic administration of FK565 induces a coronary arteritis that recapitulates histopathological and immunological features of human KD^5,9^, allowing the causal contribution of individual cell types to be investigated.

In patients with KD, a functional polymorphism in *ITPKC*, which encodes a negative regulator of calcium signaling to nuclear factor of activated T cells (NFAT), is associated with both disease susceptibility and CAA formation^10^, and a KD-associated *CASP3* variant abolishes NFAT binding^11^. Moreover, cyclosporine A (CyA), which blocks calcineurin upstream of NFAT, reduced coronary artery abnormalities when added to primary IVIG treatment in a randomized trial of patients at predicted high risk of IVIG nonresponse^12^. Yet the immune cell types through which this genetic risk operates in KD have not been established. In parallel, acute KD is associated with elevated chemokine levels^13^ that coordinate leukocyte migration and adhesion to activated endothelium. However, how genetically implicated signaling pathways and chemotactic signals combine to generate pathogenic effector cells and direct them to the coronary arteries remains unresolved.

Natural killer (NK) cells are innate lymphocytes that rapidly exert cytotoxicity through granzymes and perforin, and secrete inflammatory cytokines and chemokines^14^. Their effector capacity develops through terminal differentiation regulated by transcription factors including T-bet^15,16^. Beyond their canonical roles in antiviral and antitumor immunity, NK cells also contribute to cardiovascular injury and inflammation^17,18^, and single-cell studies of patients with KD have described a reduced but activated circulating NK cell compartment during the acute phase^19,20^. Activated NK cells upregulate the chemokine receptor CX3CR1, whose membrane-anchored ligand CX3CL1 mediates firm adhesion and migration^21^; this CX3CL1–CX3CR1 axis could therefore control site-specific NK cell accumulation. Whether NK cells contribute directly to coronary injury in KD, however, remains unknown, as do the identity of pathogenic NK cell subsets and the mechanisms directing their homing to the coronary arteries.

Here, we combined the FK565-induced mouse model of KD with comprehensive transcriptomic analysis to determine whether and how NK cells contribute to coronary arteritis. We identify NK cells as major drivers of experimental coronary arteritis, acting both as direct cytotoxic effectors and as a hub recruiting myeloid cells through the CCL5–CCR1 axis. This activity is concentrated in a terminally differentiated, T-bet-driven subset that depends on CX3CR1 for accumulation in the inflamed coronary arteries. By integrating publicly available single-cell transcriptomic data from patients, we identify a corresponding CX3CR1^+^ NK cell population that is enriched in KD. Moreover, calcineurin–NFAT signaling acts upstream of T-bet to drive this terminal differentiation. Together, these findings identify an NFAT–T-bet–CX3CR1 axis that couples terminal NK cell differentiation to site-specific vascular inflammation.

## Results

### NK cells drive coronary arteritis in FK565-treated mice

To characterize the immune cell landscape during the development of coronary arteritis, we examined the hearts of FK565-treated mice (Extended Data Fig. 1a). Histology of the coronary arteries and aortic roots revealed prominent cellular infiltration compared with control mice (Fig. 1a). Inflammation score and infiltrating cell numbers were elevated on days 5 and 7 (Fig. 1b and Extended Data Fig. 1b).

**Fig. 1:**
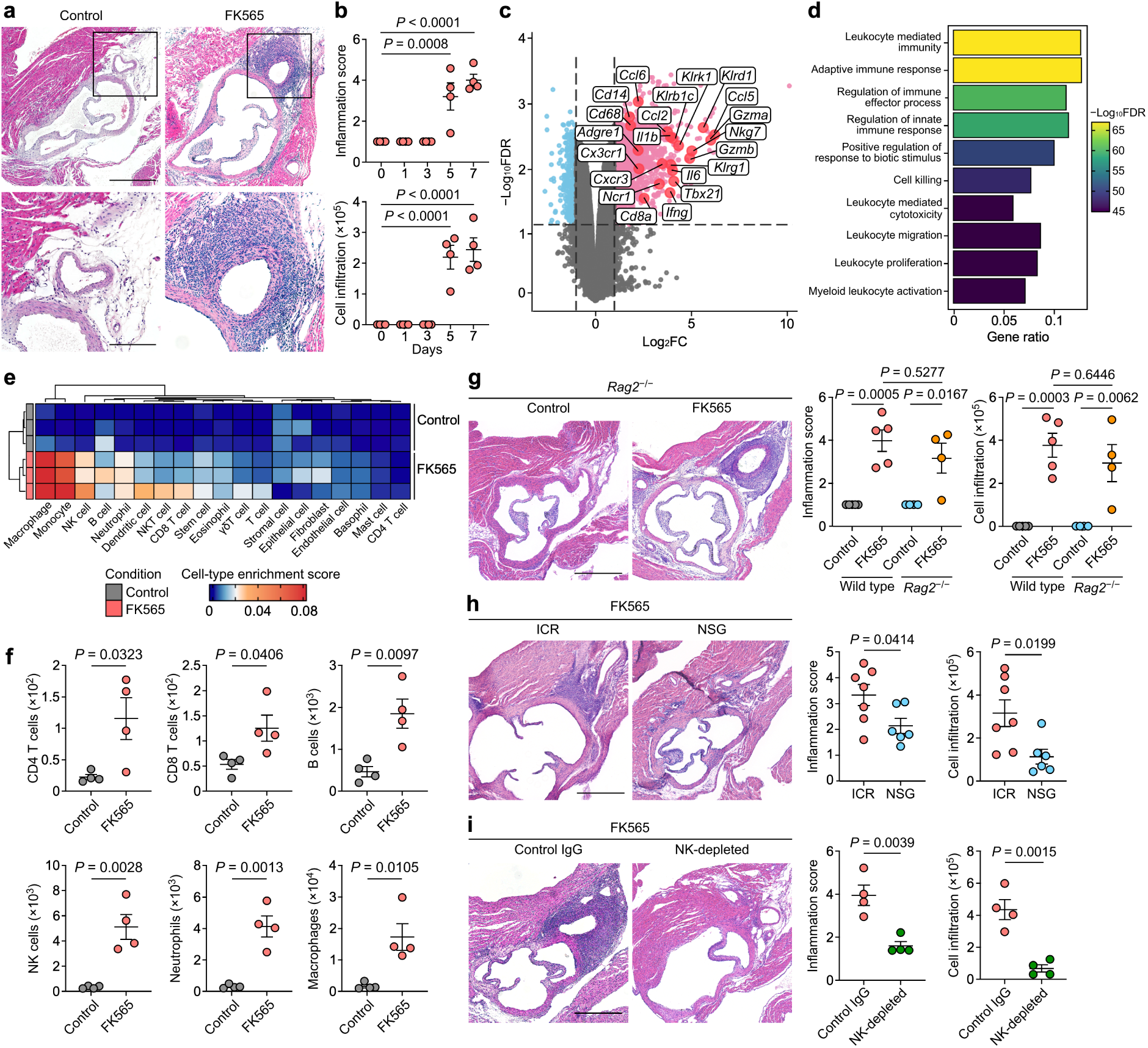
NK cells are involved in the pathogenesis of coronary arteritis in a mouse model of KD. **a**, Representative images of H&E-stained sections of the coronary arteries and the aortic roots from control and FK565-treated mice on day 7. Scale bars, 500 μm (top) and 100 μm (bottom). **b**, Inflammation score (top) and relative number of infiltrating cells per coronary artery area (bottom) in the coronary arteries of FK565-treated mice on days 0, 1, 3, 5 and 7 (*n* = 4 per time point). **c**, Volcano plot showing DEGs identified by microarray analysis of the proximal coronary arteries from control and FK565-treated mice on day 7. Genes of interest, including inflammatory and immune cell markers, are highlighted (*n* = 3). **d**, Top 10 Gene Ontology Biological Process terms enriched among upregulated genes (log_2_FC > 1, adjusted *P* < 0.05) in the proximal coronary arteries of FK565-treated mice compared with control mice. **e**, Cell-type enrichment scores determined by digital deconvolution of the microarray dataset from the proximal coronary arteries of control and FK565-treated mice. **f**, Flow cytometric quantification of immune cell populations in the heart of control and FK565-treated mice on day 7. Cell types were identified as follows: CD4 T cells (CD45^+^CD3^+^CD4^+^CD8^−^), CD8 T cells (CD45^+^CD3^+^CD4^−^CD8^+^), B cells (CD45^+^CD3^−^B220^+^), NK cells (CD45^+^CD3^−^NK1.1^+^), neutrophils (CD45^+^Ly6G^+^) and macrophages (CD45^+^Ly6G^−^F4/80^+^) (*n* = 4). **g**–**i**, Representative images of H&E-stained sections (left), inflammation score (middle) and relative number of infiltrating cells per coronary artery area (right) for the coronary arteries of (g) control or FK565-treated wild-type and *Rag2*^−/−^ mice (wild type: *n* = 5, *Rag2*^−/−^: *n* = 4), (h) FK565-treated ICR or NSG mice (ICR: *n* = 7, NSG: *n* = 6), and (i) FK565-treated mice receiving control IgG or NK-depleting antibody (*n* = 4) on day 7. Scale bars, 500 μm. Data in **b**, **f**, **g**, **h** and **i** are presented as mean ± s.e.m., with individual dots representing individual mice. Statistical significance was assessed using one-way ANOVA followed by a Dunnett multiple comparisons test for **b** or Tukey multiple comparisons test for **g**, and two-tailed unpaired *t* tests for **f**, **h** and **i**. In **c** and **d**, *P* values were adjusted for multiple testing using the Benjamini–Hochberg FDR method. Data in **b**, **f**, **g** and **i** are representative of more than three independent experiments, whereas data in **h** are pooled from two independent experiments.

To define the associated molecular signatures, we profiled the proximal coronary arteries using microarray analysis. Genes encoding cytotoxic effector molecules (*Gzmb*, *Nkg7*) and NK cell receptors (*Ncr1*, *Klrb1c*, *Klrk1*, *Klrd1*), together with *Tbx21*, *Klrg1*, *Ifng* and *Ccl5*, were among the most strongly upregulated, alongside myeloid markers (*Cd14*, *Cd68*, *Adgre1*) and the chemokine receptors *Cxcr3* and *Cx3cr1* (Fig. 1c). Gene Ontology enrichment analysis of the upregulated genes identified leukocyte-mediated immunity, cell killing and leukocyte-mediated cytotoxicity among the most enriched processes (Fig. 1d). To identify the cellular compartments involved, we deconvoluted the microarray dataset, revealing enrichment of macrophage, monocyte, NK cell and neutrophil signatures (Fig. 1e). Flow cytometric quantification revealed that numbers of CD4 T cells, CD8 T cells, B cells, NK cells, neutrophils and macrophages increased in the heart, whereas only NK cells, neutrophils and monocytes increased in the blood (Fig. 1f and Extended Data Fig. 1c, d).

The prominence of NK cell-associated transcripts led us to ask whether NK cells drive coronary arteritis. FK565-treated *Rag2*^−/−^ mice, which lack T and B cells, developed coronary arteritis comparable to wild-type mice (Fig. 1g), indicating that these cells are dispensable. By contrast, arteritis was attenuated in NSG mice, which additionally lack NK cells^22^, relative to immunocompetent ICR mice (Fig. 1h). To assess the contribution of NK cells directly, we depleted them using an antibody, which profoundly suppressed disease (Fig. 1i and Extended Data Fig. 1e). Together, these results identify NK cells as major drivers of FK565-induced coronary arteritis.

### Single-cell transcriptomics identifies NK cells as cytotoxic effectors and an immune cell hub in coronary arteritis

To resolve the cellular networks associated with coronary arteritis, we analyzed single-cell transcriptomes of immune cells from blood and hearts of control and FK565-treated mice. Clustering of 46,329 cells identified 28 populations, visualized using uniform manifold approximation and projection (UMAP; Fig. 2a and Extended Data Fig. 2a). Consistent with our microarray data, the NK cell clusters (NK_1–NK_3) highly expressed genes encoding cytotoxic effectors (*Gzma*, *Gzmb*, *Prf1*, *Fasl*) and proinflammatory mediators (*Ifng* and *Ccl5*), with the highest levels in NK_1 (Fig. 2b). Given this pronounced granzyme B expression, we asked whether NK cell-mediated cytotoxicity contributes to pathogenesis. A granzyme B inhibitor attenuated FK565-induced arteritis (Fig. 2c), supporting a pathogenic role for granzyme B. Its prominent expression in NK cells, together with disease development in the absence of T and B cells, implicates NK cells as a relevant source (Figs. 1g and 2b).

**Fig. 2:**
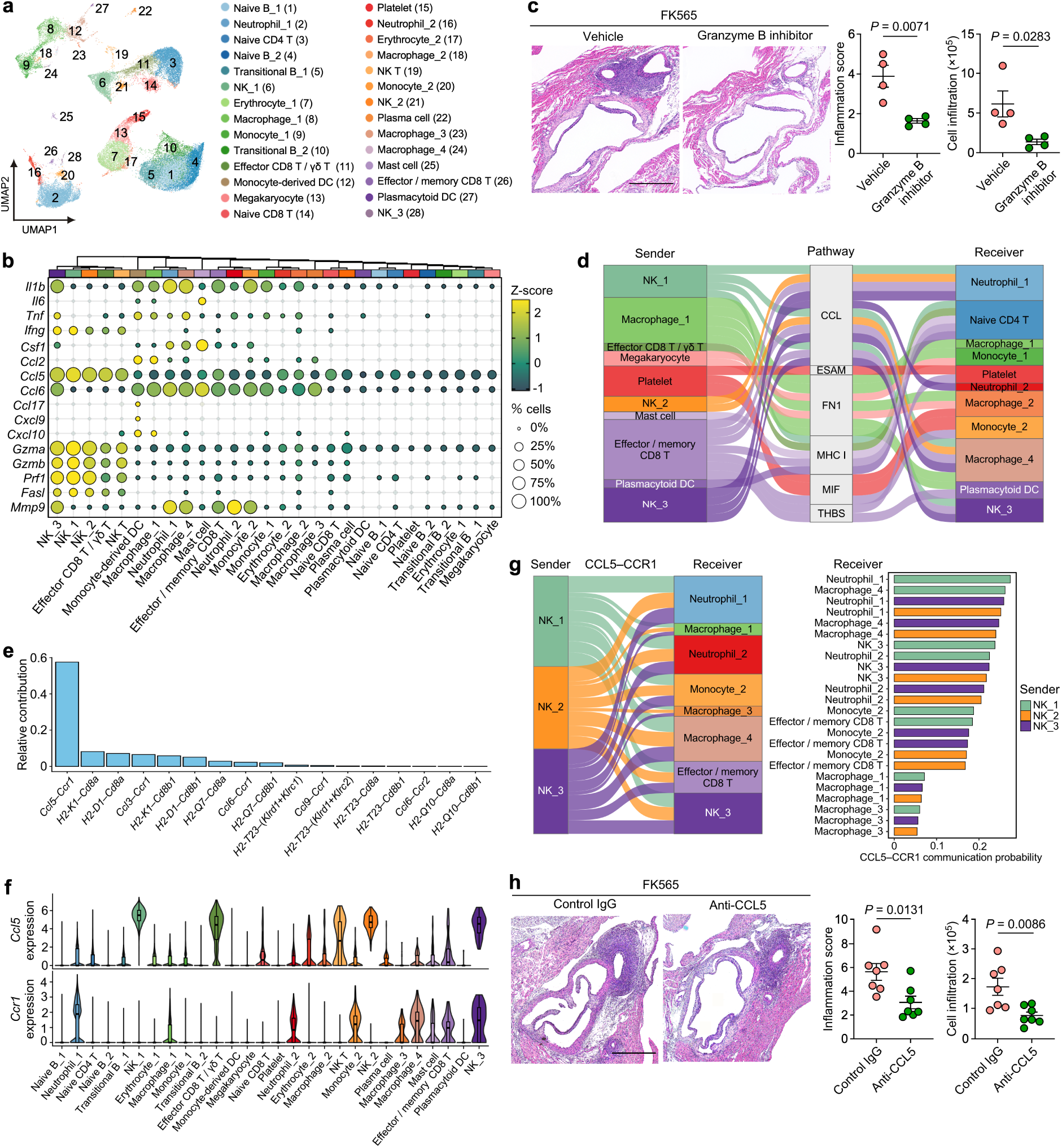
NK cell-associated cytotoxicity and effector-cell recruitment contribute to experimental coronary arteritis. **a**, UMAP embedding of cells (46,329 cells across 28 clusters) from peripheral blood and heart of control or FK565-treated mice. **b**, Dot plot showing the expression of selected genes involved in inflammation, cytotoxicity and cell recruitment across annotated clusters. **c**, Representative images of H&E-stained sections (left), inflammation score (middle) and relative number of infiltrating cells per coronary artery area (right) for the coronary arteries of FK565-treated mice receiving vehicle or granzyme B inhibitor on day 7 (*n* = 4). Scale bars, 500 μm. **d**, Alluvial plot displaying the inferred cell–cell communication network in the heart of FK565-treated mice, linking sender, signaling pathway and receiver populations. The top 30 sender–pathway–receiver combinations with the highest summed communication probability are shown. Ribbon width represents the summed communication probability. **e**, Relative contribution of individual ligand–receptor pairs to inferred NK cell-derived signaling (NK_1, NK_2 and NK_3) within the CCL and MHC I pathways in the heart of FK565-treated mice. **f**, Expression of *Ccl5* and *Ccr1* across the annotated clusters. **g**, Alluvial plot visualizing the inferred CCL5–CCR1 communication from NK cell senders (NK_1, NK_2 and NK_3) to receiver populations (left). CCL5–CCR1 communication probability for each sender–receiver pair (right). **h**, Representative images of H&E-stained sections (left), inflammation score (middle) and relative number of infiltrating cells per coronary artery area (right) for the coronary arteries of FK565-treated mice receiving control IgG or anti-CCL5 antibody on day 7 (*n* = 7). Scale bars, 500 μm. Data in **c** and **h** are presented as mean ± s.e.m., with individual dots representing individual mice. Statistical significance in **c** and **h** was assessed using two-tailed unpaired *t* tests. Data in **c** are representative of two independent experiments, whereas data in **h** are pooled from two independent experiments.

Beyond their cytotoxic potential, NK cells might also coordinate effector-cell recruitment, given the accumulation of myeloid cells, which contribute to FK565-induced coronary arteritis^23^. Cell–cell communication analysis of the FK565-treated heart inferred a network in which NK cell subsets ranked among the leading senders and signaled largely to myeloid populations (Fig. 2d and Extended Data Fig. 2b). Among NK cell-mediated outgoing signals in the CCL and MHC I pathways, *Ccl5*–*Ccr1* made the largest contribution (Fig. 2e). *Ccl5* was most highly expressed by the NK cell clusters, with lower expression in some effector T cell subsets, whereas *Ccr1* was expressed by monocytes, macrophages, neutrophils and some lymphocyte populations (Fig. 2f). Accordingly, NK cell subsets were among the strongest predicted senders of *Ccl5*–*Ccr1* signals (Extended Data Fig. 2c), with high inferred communication probabilities for neutrophils and macrophages (Fig. 2g). Systemic neutralization of CCL5 significantly reduced coronary arteritis (Fig. 2h). NK cells are thus the principal source of granzymes and CCL5 in experimental coronary arteritis, and both effector arms contribute to disease development.

### Terminally differentiated, T-bet-driven NK cells are the predominant effectors in FK565-induced coronary arteritis

To identify which NK cell population drives arteritis progression, we compared the three NK cell clusters. NK_1 was the largest and showed the highest expression of cytotoxic mediators and activation markers (Extended Data Fig. 3a, b); its subclustering identified seven subclusters (mNK1–mNK7; Fig. 3a). In control mice, blood and cardiac NK cells occupied largely distinct regions of the UMAP embedding: those from the blood consisted mainly of mNK3 and mNK4, and those from the heart almost entirely of mNK1 (Fig. 3a, b). FK565 treatment shifted both compartments into a shared region formed by mNK2 and part of mNK5, and the heart additionally gained mNK6 (Fig. 3a, b). mNK2 and mNK6 co-expressed *Gzmb*, *Ccl5*, *Klrg1* and *Tbx21* (encoding T-bet), whereas mNK5 expressed these genes only moderately but had the highest *Cd27* and *Tcf7* expression, consistent with an intermediate rather than an effector state (Fig. 3c). mNK2 was enriched for leukocyte-mediated cytotoxicity, lymphocyte migration and leukocyte adhesion to vascular endothelial cells, and mNK6 for leukocyte cell–cell adhesion, suggesting enhanced migratory potential (Fig. 3d and Extended Data Fig. 3c). Pseudobulk analysis showed elevated expression of genes associated with terminal differentiation (*Klrg1*, *Itgam*, *Tbx21*, *Zeb2*), inflammation (*Ifng*, *Ccl5*), cytotoxicity (*Gzmb*, *Prf1*, *Fcgr3*, *Ctsc*) and adhesion (*Icam1*, *Itgb2*, *Itgal*, *Cx3cr1*) in these subclusters (Extended Data Fig. 3d).

**Fig. 3:**
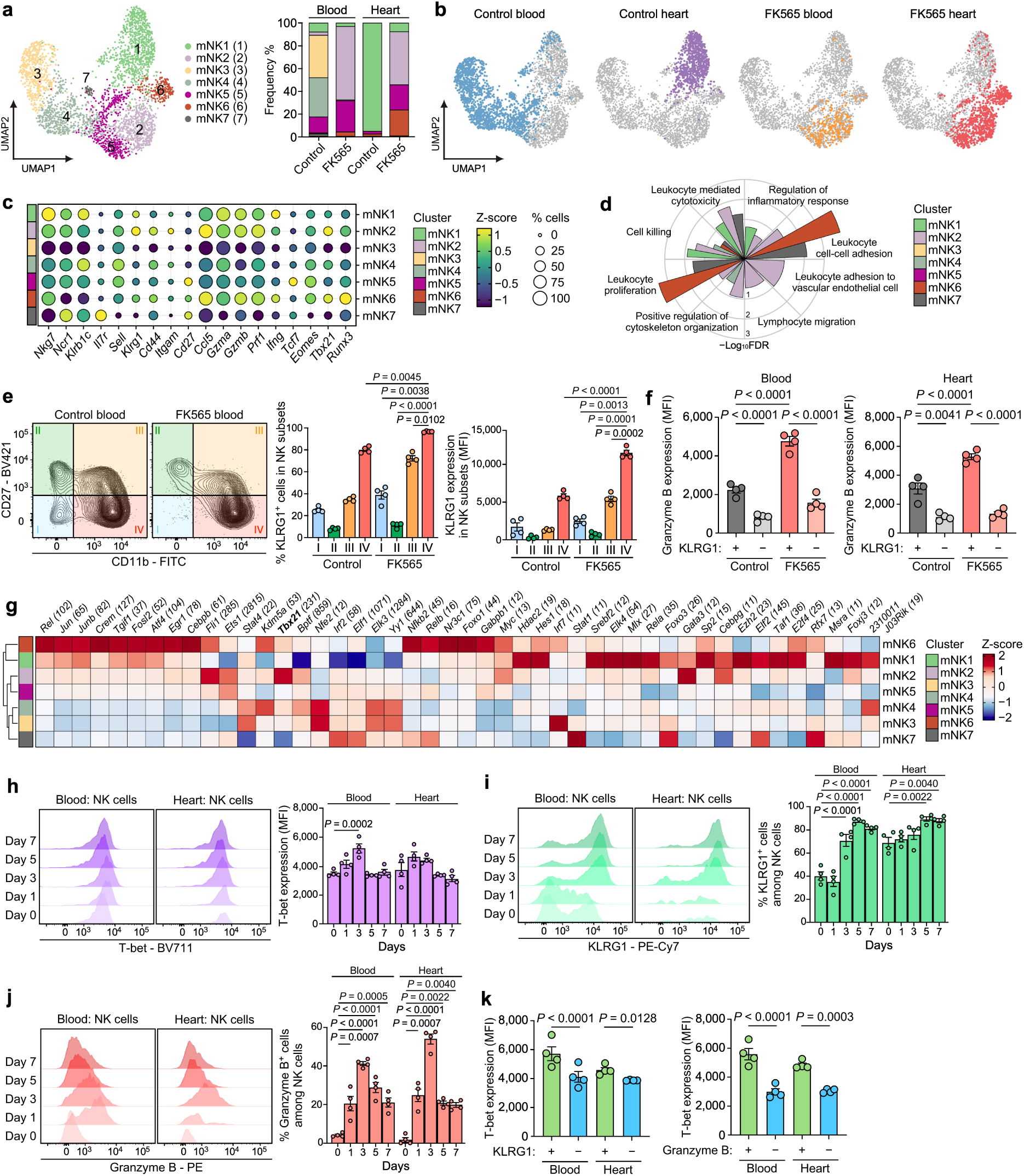
Dynamics of the NK cell subclusters expanded in FK565-treated mice. **a**, UMAP embedding of NK cell subclusters (left; 3,377 cells across seven NK subclusters) and relative frequencies of NK cell subclusters (right) from peripheral blood and heart of control and FK565-treated mice. **b**, UMAP embedding as in a, with cells from peripheral blood and heart of control and FK565-treated mice highlighted in separate plots; all other cells are shown in gray. **c**, Dot plot showing the expression of NK cell-related genes across the defined NK subclusters. **d**, Circular bar plot showing the enrichment of selected Gene Ontology Biological Process terms for significantly upregulated DEGs (log_2_FC > 0.5, adjusted *P* < 0.05) across the identified NK subclusters. **e**, Representative flow cytometry plots (left) and the frequency of KLRG1^+^ cells and KLRG1 expression in NK cell maturation subsets, based on the surface expression of CD11b and CD27, from the peripheral blood of control and FK565-treated mice (*n* = 4). **f**, Granzyme B expression in KLRG1^+^ or KLRG1^−^ NK cells from peripheral blood and heart of control and FK565-treated mice (*n* = 4). **g**, Heatmap visualizing the average inferred regulon activity scores of defined transcription factors across the identified NK subclusters in peripheral blood and heart from control or FK565-treated mice. Regulon activities were computed as area under the curve (AUC) scores at the single-cell level, averaged per subcluster and subsequently row-scaled (Z-score). The number in parentheses next to each transcription factor name indicates the number of high-confidence target genes constituting that specific regulon. **h**, Representative flow cytometry histograms and the corresponding expression of T-bet in NK cells isolated from the blood and heart of FK565-treated mice at the indicated time points (*n* = 4). **i** and **j**, Representative flow cytometry histograms and the corresponding frequency of KLRG1^+^ (i) and granzyme B^+^ (j) cells among NK cells isolated from the blood and heart of FK565-treated mice at the indicated time points (*n* = 4). **k**, Expression levels (mean fluorescence intensity, MFI) of T-bet in KLRG1^+^ or KLRG1^−^, and granzyme B^+^ or granzyme B^−^ NK cells from the peripheral blood or the heart of FK565-treated mice (*n* = 4). Data in **e**, **f** and **h**–**k** are presented as mean ± s.e.m., with individual dots representing individual mice. Statistical significance was assessed using two-way ANOVA with repeated measures on maturation stage and the Geisser–Greenhouse correction followed by a Šídák multiple comparisons test for **e**, and two-way ANOVA with repeated measures on KLRG1 status followed by a Šídák multiple comparisons test for **f**. For **h**–**j**, ordinary one-way ANOVA with a Dunnett multiple comparisons test was used. Statistical significance in **k** was assessed using two-tailed ratio paired *t* tests. Data in **e**, **f** and **h**–**k** are representative of three independent experiments.

We next assessed NK cell maturation across four stages defined by CD11b and CD27 expression^24^. The proportions of cells in stages I and IV differed between groups, with no significant differences in stages II and III (Extended Data Fig. 3e). In terminally differentiated stage IV, both the frequency of KLRG1^+^ cells and KLRG1 expression were increased in FK565-treated mice (Fig. 3e), consistent with the expansion of mNK2, which had low *Cd27* and the highest *Itgam* (encoding CD11b) and *Klrg1* expression (Fig. 3c). These KLRG1^+^ NK cells expressed more granzyme B than their KLRG1^−^ counterparts, and more still in FK565-treated mice (Fig. 3f and Extended Data Fig. 3f).

The *Tbx21* regulon showed high inferred activity in mNK2, with weaker positive activity in mNK6 (Fig. 3g), and T-bet expression in blood NK cells peaked on day 3 (Fig. 3h). Reanalysis of published RNA-seq data from *Tbx21*^−/−^ NK cells^25^ showed lower expression of *Gzmb* and *Ccl5* than in wild-type cells (Extended Data Fig. 3g), consistent with the established role of T-bet in terminal maturation^15^. The proportion of KLRG1^+^ NK cells increased from day 3 in the blood and day 5 in the heart (Fig. 3i), and that of granzyme B^+^ NK cells peaked on day 3 and remained elevated through day 7 (Fig. 3j). T-bet expression was higher in KLRG1^+^ and granzyme B^+^ NK cells than in their negative counterparts (Fig. 3k). The cytotoxic and inflammatory program is thus concentrated in a terminally differentiated, T-bet-driven NK cell subset that expands in FK565-induced coronary arteritis.

### A terminally differentiated NK cell state, pathogenic in FK565-induced coronary arteritis, is shared with human KD

To examine the relevance of our findings to human disease, we analyzed publicly available single-cell transcriptomic datasets of peripheral blood mononuclear cells from patients with KD and healthy controls^19,26,27,28^. Subclustering the NK cell cluster (Extended Data Fig. 4a, b) resolved 18,336 NK cells into six subclusters (hNK1–hNK6), of which hNK1 accounted for a greater proportion of NK cells in KD datasets (Fig. 4a). hNK1 highly expressed *KLRG1*, *B3GAT1* (encoding the CD57-synthesizing enzyme), *CCL5*, *GZMB* and *TBX21,* with low *IL7R*, *SELL* and *TCF7*, consistent with a terminally differentiated, cytotoxic phenotype (Fig. 4b). This subcluster was enriched for gene programs associated with cell killing, leukocyte-mediated cytotoxicity and leukocyte cell–cell adhesion, paralleling mNK2 (Fig. 4c).

**Fig. 4:**
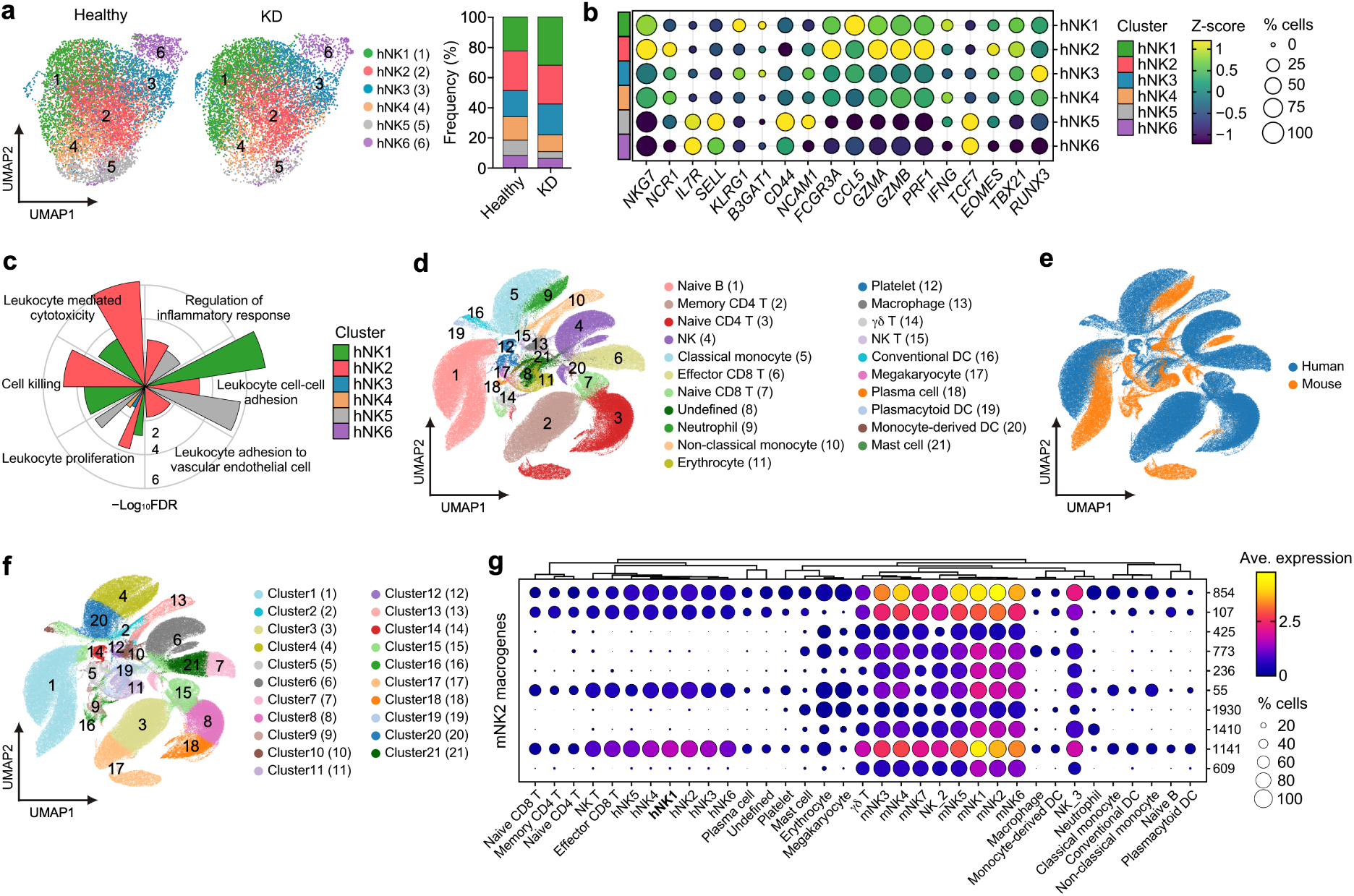
Cross-species integration identifies a shared T-bet^+^ effector NK cell state in human KD and the FK565 mouse model. **a**, UMAP embedding of human NK cell subclusters (left; 18,336 cells across six NK subclusters) and their relative frequencies (right) from peripheral blood of healthy donors and patients with KD. **b**, Dot plot showing the expression of human NK cell-related genes across the defined human NK subclusters. **c**, Circular bar plot showing the enrichment of selected Gene Ontology Biological Process terms for significantly upregulated DEGs (log_2_FC > 0.5, adjusted *P* < 0.05) across the identified human NK subclusters. **d**, UMAP embedding of the SATURN cross-species integration of human and mouse immune cells, colored by the 21 coarse cell-type labels used for integration. **e**, Species identity overlaid on the SATURN UMAP. **f**, UMAP colored by K-means clusters (k = 21) computed on the SATURN embedding. **g**, Dot plot showing the expression of the top 10 SATURN macrogenes (groups of functionally related genes) most enriched in mNK2 across human and mouse immune cell populations, including the identified human and mouse NK cell subclusters. The 10 highest-weighted genes of the top 10 macrogenes and the plotted values are provided in Supplementary Tables 1 and 2.

To compare NK cell states across species, we integrated the human and mouse datasets into a shared space generated by SATURN^29^. Cells were organized by cell type rather than by species (Fig. 4d, e and Extended Data Fig. 4c), and K-means clustering placed human and mouse NK cells in shared clusters (Fig. 4f), indicating that, at least for the NK cell population, the gene expression profiles were sufficiently conserved to allow meaningful cross-species comparison.

mNK2-enriched SATURN macrogenes were expressed in human NK subclusters, more highly than in other human immune populations, with relatively high levels in hNK1 (Fig. 4g and Supplementary Tables 1 and 2). Consistently, mNK2 showed its highest transcriptome-wide correlation with hNK1, and the signature of each scored highest in the other (Extended Data Fig. 4d–f). By contrast, the mNK6 signature scored highest in hNK5 (Extended Data Fig. 4g), an immature subset with high *NCAM1* (encoding CD56), low *FCGR3A* (encoding CD16) and limited cytotoxic gene expression (Fig. 4b). Whether a human counterpart of the predominantly cardiac mNK6 population exists remains unresolved because the human datasets sampled peripheral blood. These results establish that the pathogenic NK cell state expanding in FK565-treated mice corresponds to a population enriched in patients with KD.

### CX3CR1 directs pathogenic NK cells to the coronary arteries and promotes disease development in mice

To understand how the shared NK cell state localized to the coronary arteries, we compared functional gene programs across human and mouse subclusters. In hNK1 and mNK2, signatures of cytotoxicity, integrin-mediated adhesion and effector transcription factors were broadly shared and accentuated in KD and FK565-treated mice (Extended Data Fig. 5a, b). Among chemokine receptor genes, *CX3CR1* and *Cx3cr1* were consistently enriched in hNK1 and mNK2, respectively, and further upregulated under disease conditions (Fig. 5a). In FK565-treated mice, the proportion of CX3CR1^+^ NK cells increased progressively in blood and heart (Fig. 5b), and these cells expressed higher levels of KLRG1, granzyme B and T-bet than their CX3CR1^−^ counterparts (Fig. 5c). Consistent with a role for T-bet in driving this phenotype, *Tbx21*-deficient NK cells expressed less *Cx3cr1* than wild-type NK cells (Extended Data Fig. 3g).

**Fig. 5:**
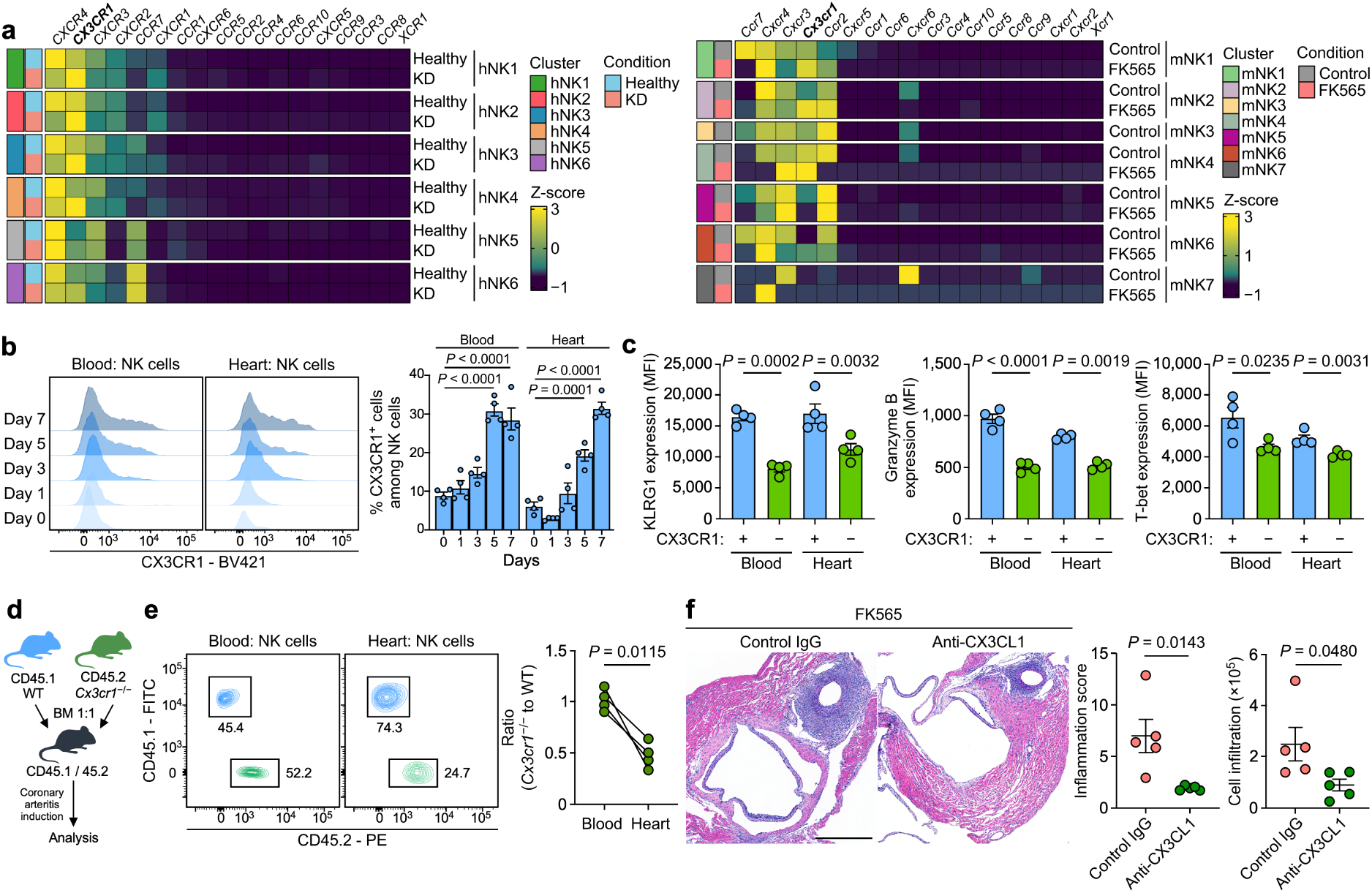
The CX3CL1–CX3CR1 axis promotes cardiac NK cell accumulation and coronary arteritis in FK565-treated mice. **a**, Heatmap visualizing the scaled average expression of chemokine receptor genes across the identified human NK cell subclusters (left) and mouse NK cell subclusters (right). Average gene expression levels were calculated for each condition (healthy versus KD and control versus FK565) within each respective subcluster and subsequently scaled to compute Z-scores. **b**, Representative flow cytometry histograms and the corresponding frequency of CX3CR1^+^ cells among NK cells isolated from the blood and heart of FK565-treated mice at the indicated time points (*n* = 4). **c**, Expression levels (MFI) of KLRG1, granzyme B and T-bet in CX3CR1^+^ and CX3CR1^−^ NK cells from the peripheral blood and the heart of FK565-treated mice (*n* = 4). **d**, Schematic illustration of the protocol: bone marrow (BM) cells from CD45.1 wild-type and CD45.2 *Cx3cr1*^−/−^ mice were adoptively transferred into irradiated CD45.1/CD45.2 heterozygous recipient mice. After 7 weeks, coronary arteritis was induced. **e**, Representative flow cytometry plot of CD45.1 (wild type) against CD45.2 (*Cx3cr1*^−/−^) gated on NK cells (left). The ratio of *Cx3cr1*^−/−^ NK cells to wild-type NK cells in the peripheral blood and the hearts of FK565-treated mixed bone marrow chimeric mice (right) (*n* = 4). **f**, Representative images of H&E-stained sections (left), inflammation score (middle) and relative number of infiltrating cells per coronary artery area (right) for the coronary arteries of FK565-treated mice receiving control IgG or anti-CX3CL1 antibody on day 7 (*n* = 5). Scale bars, 500 μm. Data in **b, c** and **f** are presented as mean ± s.e.m., with individual dots representing individual mice. Statistical significance in **b** was determined using ordinary one-way ANOVA with a Dunnett multiple comparisons test. Statistical significance in **c**, **e** and **f** was assessed using two-tailed ratio paired *t* tests for **c**, a two-tailed paired *t* test for **e** and a two-tailed unpaired *t* test for **f**. Data in **b**, **c** and **f** are representative of three independent experiments. Data in **e** are representative of two independent experiments.

To test whether CX3CR1 is required for cardiac accumulation of NK cells, we generated mixed bone marrow chimeras reconstituted with wild-type (CD45.1) and *Cx3cr1*^−/−^ (CD45.2) cells at a 1:1 ratio (Fig. 5d and Extended Data Fig. 5c). *Cx3cr1*^−/−^ NK cells were selectively underrepresented in the heart relative to the blood (Fig. 5e and Extended Data Fig. 5d, e), demonstrating a cell-intrinsic requirement for CX3CR1 in cardiac NK cell accumulation.

We next asked which cells in the coronary artery express the CX3CR1 ligand, CX3CL1. In human coronary artery and endothelial cell atlases^30,31^, *CX3CL1* expression was largely restricted to endothelial cells (Extended Data Fig. 6a–d). CX3CL1 was likewise detected along the luminal surface of coronary arteries in both control and FK565-treated mice, without a significant increase in *Cx3cl1* expression (Extended Data Fig. 6e, f). A neutralizing anti-CX3CL1 antibody significantly attenuated FK565-induced coronary arteritis (Fig. 5f). CX3CR1 thus contributes to cardiac NK cell accumulation and CX3CL1 to disease severity in FK565-treated mice, with corresponding receptor and ligand expression in human datasets.

### NFAT signaling accompanies the differentiation of pathogenic NK cells

KD is genetically linked to NFAT signaling through a functional *ITPKC* polymorphism^10^. We therefore examined the relationship of NFAT signaling to the differentiation of pathogenic NK cells. RNA velocity and pseudotime analyses inferred a differentiation trajectory from the immature subclusters (mNK3 and mNK4) through mNK5 toward the pathogenic effector subclusters mNK2 and mNK6 in FK565-treated mice (Fig. 6a, b). Along this inferred trajectory, expression of *Nfatc1*, *Nfatc2*, *Nfatc3* and the calcineurin catalytic subunits *Ppp3ca*, *Ppp3cb* and *Ppp3cc* increased earlier in pseudotime than *Tbx21*, whose increase preceded or coincided with increases in *Klrg1*, *Gzmb*, *Ccl5* and *Cx3cr1* (Fig. 6a, b). The AP-1 component *Jun*, a transcriptional partner of NFAT^32^, was also induced concurrently with *Tbx21* along the trajectory (Fig. 6a, b). mNK2 and mNK6 highly expressed calcineurin–NFAT pathway components and were enriched for NFAT- and calcineurin-associated gene programs (Fig. 6c, d). Among the NFAT family members examined, *Nfatc1* most strongly correlated with *Tbx21* within mNK2 metacells (Fig. 6e).

**Fig. 6:**
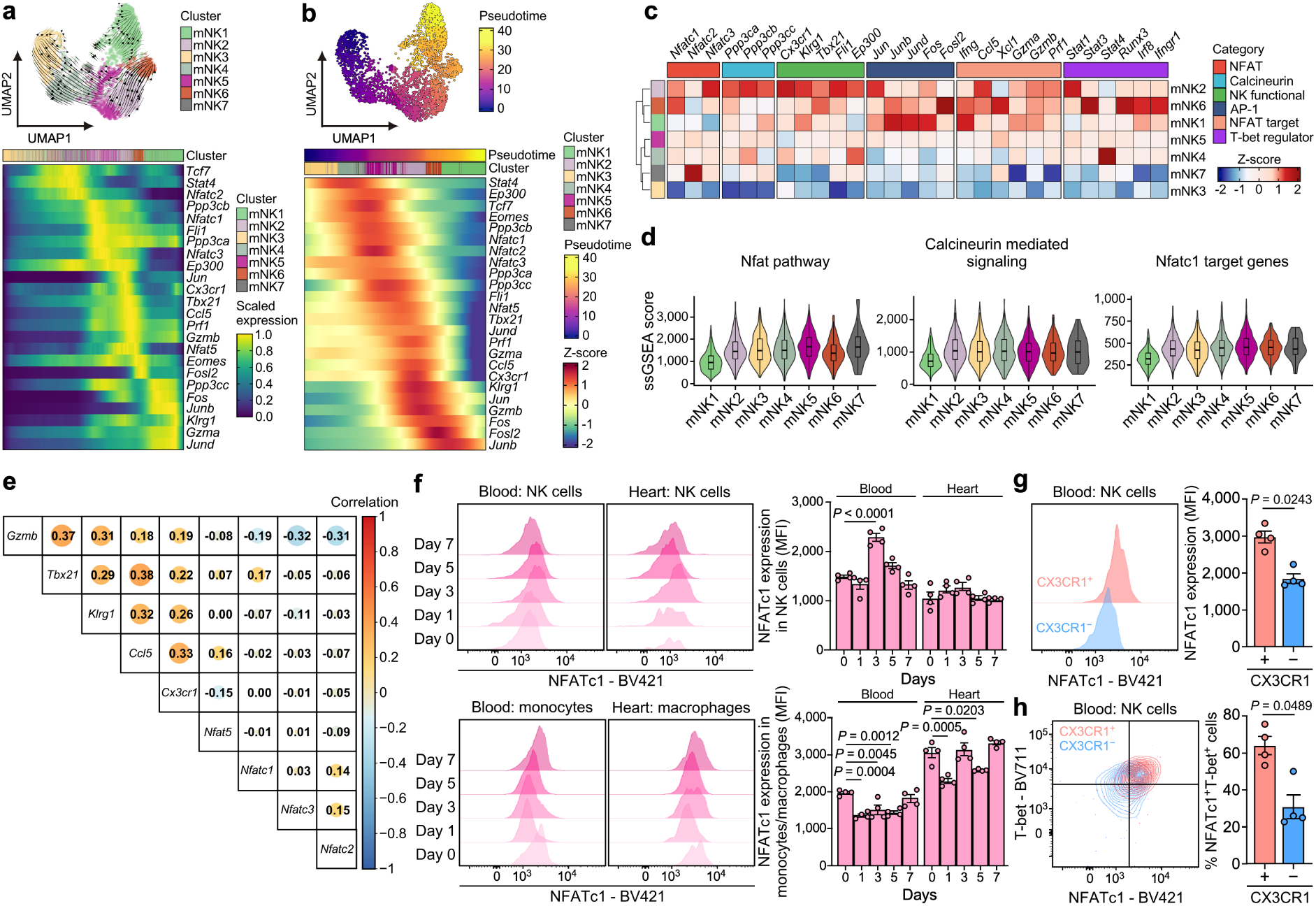
NFATc1 is coordinately induced with T-bet in pathogenic NK cells. **a**, **b**, Trajectory inference of identified mNK subclusters. mNK subclusters from peripheral blood and heart of control and FK565-treated mice were analyzed using RNA velocity-based scVelo (a) and pseudotime inference with Monocle3 (b). UMAP projections overlaid with RNA velocity streamlines (a) or pseudotime scores (b), illustrating inferred transitions among NK cell subclusters (top). The expression dynamics of NK cell- and NFAT-related genes along the inferred trajectories (bottom). **c**, Heatmap visualizing the pseudobulk gene expression of selected markers for the calcineurin–NFAT pathway, AP-1 transcription factor complex and NK cell functional regulators across mNK cell subclusters (mNK1–mNK7). Gene expression levels were aggregated per subcluster and subsequently scaled per gene to compute Z-scores. **d**, Violin plots illustrating the distribution of ssGSEA scores for the indicated functional modules across mNK cell subclusters. **e**, Pairwise Spearman correlation coefficients between the indicated genes across mNK2 metacells, constructed using hdWGCNA (k = 25 nearest neighbors). Circle size and color indicate the correlation coefficient. **f**, Representative flow cytometry histograms and the corresponding expression of NFATc1 in blood and cardiac NK cells, blood monocytes and cardiac macrophages of FK565-treated mice at the indicated time points (*n* = 4). **g**, Representative flow cytometry histograms (left) and the corresponding quantification of NFATc1 expression (right) in CX3CR1^+^ and CX3CR1^−^ NK cells isolated from the blood of FK565-treated mice on day 3 (*n* = 4). **h**, Representative density plots showing the co-expression of NFATc1 with T-bet in CX3CR1^+^ and CX3CR1^−^ NK cells from the peripheral blood of FK565-treated mice on day 3 (left). Quantification of the percentage of NFATc1^+^T-bet^+^ cells within CX3CR1^+^ and CX3CR1^−^ NK cell populations (right) (*n* = 4). Data in **f**, **g** and **h** are presented as mean ± s.e.m., with individual dots representing individual mice. Statistical significance in **g** and **h** was assessed using a two-tailed ratio paired *t* test for **g** and a two-tailed paired *t* test for **h**. In **f**, statistical significance was assessed using ordinary one-way ANOVA with a Dunnett multiple comparisons test. Data in **f**, **g** and **h** are representative of three independent experiments.

Comparable relationships were evident in NK cell transcriptomic data from patients with KD. *NFATC3* and *PPP3CB* expression increased earlier in pseudotime than *TBX21* and the effector genes along the human NK cell trajectory (Extended Data Fig. 7a). Calcineurin– NFAT pathway components were most highly expressed in hNK3, immediately upstream of hNK1 along the trajectory (Extended Data Fig. 7a, b), and within hNK1 metacells, *NFATC3* correlated most strongly with *TBX21*, *KLRG1* and *CX3CR1* (Extended Data Fig. 7c). The calcineurin–NFAT program thus accompanies the T-bet effector state in both species, although the associated NFAT family member differs.

NFATc1 protein expression increased in parallel with T-bet in NK cells, peaking on day 3 in the blood, but not in blood monocytes or cardiac macrophages (Fig. 6f), which are themselves required for arteritis in this model^23^. CX3CR1^+^ NK cells expressed more NFATc1 than their CX3CR1^−^ counterparts (Fig. 6g) and were enriched for cells co-expressing NFATc1 and T-bet (Fig. 6h), linking their coordinated expression to the terminally differentiated, CX3CR1^+^ effector state.

To explore a direct regulatory link, we examined published chromatin datasets. The regulatory element in the *Tbx21* locus (conserved non-coding sequence 12; *CNS12*)^16^ containing NFAT, STAT4 and AP-1 motifs became accessible and was occupied by STAT4 and p300 upon NK cell activation; the sequences and spacing of these motifs were conserved between mouse and human (Extended Data Fig. 8a, b). Collectively, these results identify NFATc1 and T-bet as coordinately expressed during the development of the pathogenic state, with *Nfatc1* expression increasing earlier than *Tbx21* along the inferred trajectory.

### Calcineurin inhibition suppresses pathogenic NK cells and coronary arteritis in FK565-treated mice

To assess the contribution of calcineurin–NFAT signaling to disease development, we administered CyA to FK565-treated mice. CyA markedly suppressed coronary arteritis, reducing the inflammation score and infiltrating cell numbers (Fig. 7a). CyA also reduced disease in *Rag2*^−/−^ mice (Fig. 7b), demonstrating that its protective effect does not require T or B cells.

**Fig. 7:**
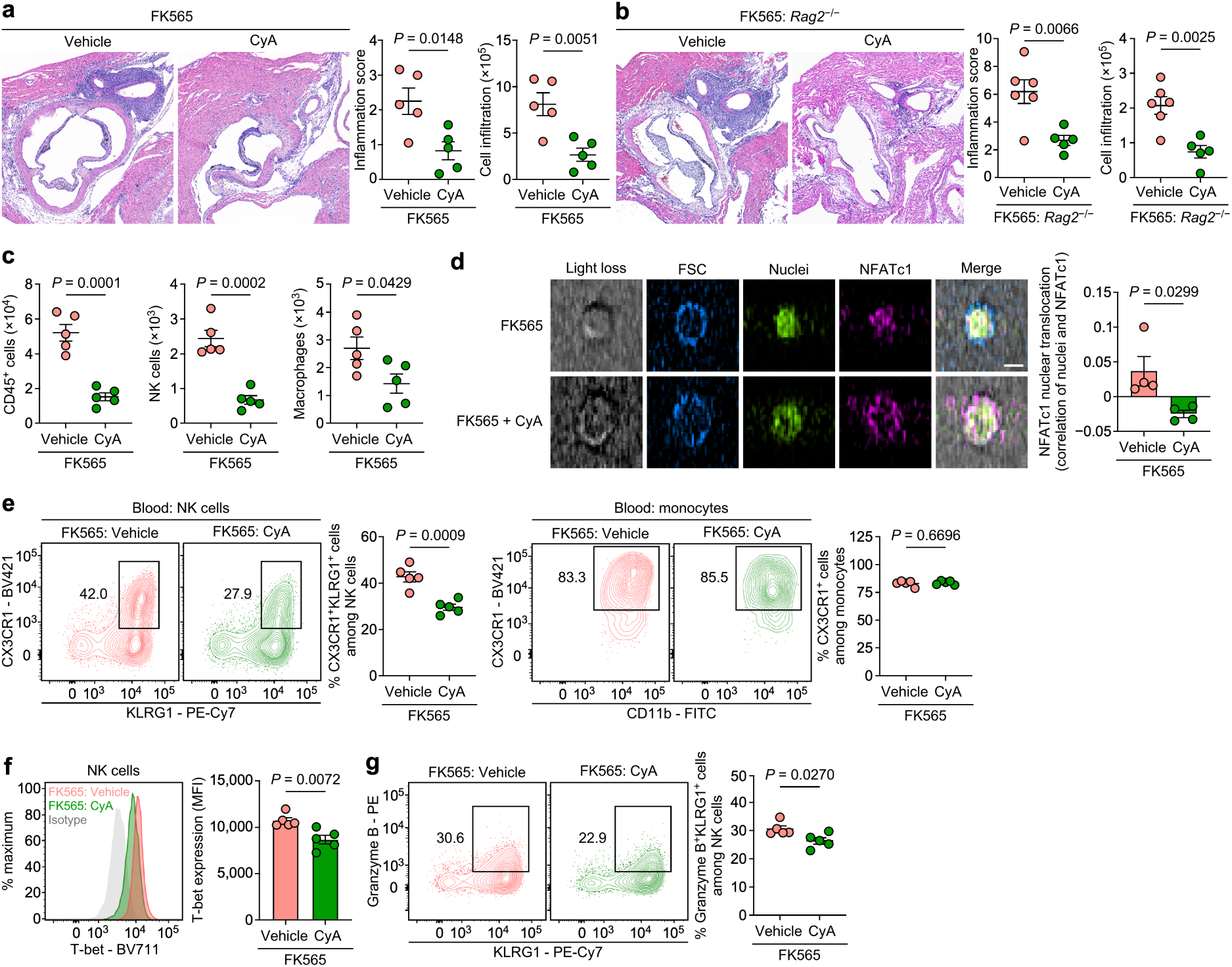
Calcineurin inhibition suppresses the pathogenic NK cell state and coronary arteritis in FK565-treated mice. **a**, **b**, Representative images of H&E-stained sections (left), inflammation score (middle) and relative number of infiltrating cells per coronary artery area (right) for the coronary arteries in wild-type (a; *n* = 5) or *Rag2*^−/−^ (b; vehicle: *n* = 6, CyA: *n* = 5) FK565-treated mice given either vehicle or cyclosporine A (CyA) on day 7. Scale bars, 500 μm. **c**, Flow cytometric quantification of CD45^+^ cells, NK cells and macrophages in the heart of FK565-treated mice receiving vehicle or CyA (*n* = 5). **d**, Representative imaging flow cytometry images of NK cells from the blood of FK565-treated mice on day 3, given vehicle or CyA, showing bright-field (light loss), forward scatter (FSC), nuclei, NFATc1 and the merged image (left), with quantification of NFATc1 nuclear translocation, measured as the correlation between the nuclear and NFATc1 images (right) (*n* = 4). Scale bar, 5 μm. **e**, Representative flow cytometry plots and the corresponding frequency of CX3CR1^+^KLRG1^+^ cells among NK cells (left) and CX3CR1^+^ cells among monocytes (right) isolated from the blood of FK565-treated mice receiving vehicle or CyA on day 7 (*n* = 5). **f**, Representative histogram and expression of T-bet in NK cells isolated from the blood of FK565-treated mice receiving vehicle or CyA on day 3 (*n* = 5). **g**, Representative flow cytometry plots and the corresponding frequency of granzyme B^+^KLRG1^+^ cells among NK cells isolated from the blood of FK565-treated mice receiving vehicle or CyA on day 7 (*n* = 5). Data in **a**–**g** are presented as mean ± s.e.m., with individual dots representing individual mice. Statistical significance in **a**–**g** was assessed using two-tailed unpaired *t* tests. Data in **a–c** and **e**–**g** are representative of three independent experiments, and data in **d** are representative of two independent experiments.

Flow cytometric analysis showed that CyA reduced the numbers of cardiac CD45^+^ cells, NK cells and macrophages (Fig. 7c), and blocked the nuclear translocation of NFATc1 in NK cells from FK565-treated mice (Fig. 7d). CyA decreased the frequency of CX3CR1^+^KLRG1^+^ NK cells, whereas that of CX3CR1^+^ monocytes was unchanged (Fig. 7e). Notably, CyA suppressed T-bet expression in NK cells on day 3 (Fig. 7f) and reduced the frequency of granzyme B^+^KLRG1^+^ NK cells (Fig. 7g), consistent with the T-bet-dependent expression of effector genes (Extended Data Fig. 3g). Together with the trajectory and chromatin analyses, these data support an upstream role for calcineurin–NFAT signaling in the T-bet-driven effector program (Fig. 6 and Extended Data Fig. 8). The reduction in cardiac macrophages (Fig. 7c) likely reflects diminished NK cell-mediated recruitment through the CCL5–CCR1 axis (Fig. 2). Collectively, these findings show that calcineurin–NFAT signaling promotes the differentiation of CX3CR1^+^ pathogenic NK cells and the development of coronary arteritis in FK565-treated mice, with a corresponding NK cell population enriched for a calcineurin– NFAT signature in KD datasets.

## Discussion

Identifying the immune cells that orchestrate arteritis at a given vascular site remains a challenge in immunology, including in KD. Combining an FK565-induced mouse model with publicly available single-cell transcriptomes from patients, we establish NK cells as major drivers of experimental coronary arteritis and characterize a terminally differentiated, T-bet-driven CX3CR1^+^ state regulated by the calcineurin–NFAT pathway, with a corresponding population enriched in human KD datasets. These findings identify a pathogenic NK cell state in coronary arteritis, extending work focused largely on myeloid and T cell biology.

In prior studies, myeloid cells, particularly CD11c^+^ macrophages, were emphasized in FK565-induced coronary arteritis^23^. We assign NK cells a role in cytotoxicity, reported after myocardial infarction^18^, and in CCL5-mediated myeloid recruitment. Consistent with NK cell involvement in human KD, NK cells from patients upregulate cytotoxic and chemokine genes including *CCL5*^20^, and the CD16^hi^ NK cell fraction declines after IVIG treatment^19^. Although granzyme B inhibition and CCL5 neutralization are systemic rather than NK cell-specific, the combined findings indicate that NK cells contribute to disease development through both mechanisms.

NK cell accumulation at the vascular wall depends on CX3CR1, whose increased expression would favor engagement of constitutively expressed endothelial CX3CL1. Whether the corresponding human subset accumulates in the coronary wall remains unresolved because affected tissue is rarely available in KD. Beyond chemoattraction and adhesion^21^, CX3CL1 enhances NK cell cytotoxicity^33^, and could therefore contribute to effector function as well as recruitment. Because the FK565 model reproduces acute arteritis but not aneurysm formation, whether NK cells also contribute to the CAAs associated with long-term morbidity remains an open question.

The genetic architecture of KD implicates calcium–NFAT signaling through a functional *ITPKC* polymorphism^10^ and related pathway variants^11^, yet these associations have been interpreted largely in the context of T cells and, for *ITPKC*, myeloid inflammasome activation^34^. We instead found that NFATc1 was induced in NK cells and coordinately expressed with T-bet along the inferred differentiation trajectory toward the pathogenic state. A potential mechanism is suggested by *CNS12* in the *Tbx21* locus, which contains NFAT- and STAT4-binding motifs, becomes accessible and is occupied by STAT4 and p300 upon NK cell activation. This element is dispensable for T-bet induction during NK cell development but IL-12–STAT4-responsive in activated NK cells^16^. In NK cells, STAT4 opens de novo enhancers upon acute activation^35,36^, providing one route by which IL-12 induces T-bet and its effector program. Such enhancers are marked and maintained by p300^37^, a histone acetyltransferase also recruited by NFAT^38^. Calcium–calcineurin and IL-12 signals may therefore converge on *CNS12* through NFAT and STAT4, recruiting p300 to promote chromatin accessibility and *Tbx21* transcription. The AP-1 component *Jun* was also induced concurrently with *Tbx21* along the inferred trajectory, and *CNS12* contains an AP-1-binding motif, raising the possibility of a further AP-1 input at this element. Whereas STAT4 and p300 occupancy was measured directly in NK cells, NFAT binding is inferred from motif analysis and occupancy in other lymphocyte lineages and awaits direct validation.

A species difference emerged in which NFAT family member accompanied the T-bet program: *Nfatc1* in mouse and *NFATC3* in human NK cells. NFAT family members share a highly conserved DNA-binding domain and recognize a common core sequence^39^, identical in the mouse and human *CNS12* elements. Both paralogs are expressed in NK cells, suggesting that the distinction reflects which member is most strongly co-regulated with the T-bet program; whether both act at the *TBX21* locus remains untested. Calcineurin inhibition acts upstream of all calcineurin-regulated NFAT members (NFATc1–NFATc4)^39^ and could therefore suppress NFAT-dependent activity mediated by either paralog. These findings provide a rationale for investigating pathway-level inhibition, although the relative benefits of targeting calcineurin or individual family members remain to be established.

Consistent with the clinical relevance of calcineurin inhibition, the addition of CyA to primary IVIG reduced coronary artery abnormalities in patients at predicted high risk of IVIG nonresponse in the randomized KAICA trial^12^. An earlier report that calcineurin inhibitors exacerbate NOD1-mediated coronary arteritis^40^ used substantially higher doses and attributed the effect to a MyD88-dependent mechanism in vascular and monocytic cells, suggesting that the effects may depend on dose and cellular target. Although calcineurin inhibition is not cell-type selective, several observations point to NK cells as a relevant target: NFATc1 was induced in NK cells but not in blood monocytes or cardiac macrophages, CyA blocked its nuclear translocation, and protection persisted in the absence of T and B cells. Systemic treatment nonetheless does not exclude effects on myeloid or vascular cells. NK cell-specific deletion of *Nfatc1*, together with direct perturbation of *NFATC3* in human NK cells, will be required to establish cell-intrinsic NFAT dependence and to test whether the same axis operates in patients.

An unresolved question is how the pathogenic NK cell program is initiated. In multisystem inflammatory syndrome in children (MIS-C), a Kawasaki-like vasculitis following SARS-CoV-2 infection^41,42^, CX3CR1^+^ cytotoxic lymphocytes that home to the vascular endothelium are activated during disease and this activation resolves with recovery^43,44^. CD16^hi^ NK cells in MIS-C, but not in pediatric COVID-19, are also enriched for a cytotoxicity gene signature^45^, paralleling the effector phenotype we identify here. Microbial stimuli activate NK cells largely indirectly, through IL-15, IL-12 and IL-18 stimulation^46,47^. Because these cytokines also converge on T-bet^15,16^, defining their interaction with the *ITPKC* risk genotype could help explain how infectious triggers contribute to KD in genetically susceptible children.

Within pathogenic NK cell subsets, regulons for *Fli1* and *Crem*, both negative regulators of NK cell function induced downstream of activation and IL-15 signaling^48,49,50^, were also prominent. Their high inferred activity may reflect a counter-regulatory response to intense activation that restrains NK cell-mediated inflammation or contributes to disease resolution. In summary, our study identifies a pathogenic NK cell state that contributes to coronary arteritis and implicates calcineurin–NFAT signaling in this state, linking a T-bet-driven, terminally differentiated effector program to CX3CR1-dependent cardiac accumulation; a corresponding NK cell population is enriched in human KD transcriptomic datasets, providing a basis for investigating its contribution to coronary inflammation in humans.

## Methods

### Mice

C57BL/6NCrSlc and ICR mice were obtained from Japan SLC. NSG (NOD.Cg-*Prkdc^scid^Il2rg^tm1Wjl^*/SzJ) mice were obtained from The Jackson Laboratory. *Cx3cr1*^fl/fl^ mice on a C57BL/6 background were generated by Advanced Animal Model Support (AdAMS), with loxP sites inserted into the intron 5′ of exon 1 and into the 3′ untranslated region of *Cx3cr1*. *Rag2*^−/−^ mice on a C57BL/6 background were kindly provided by Hiroshi Kawamoto (Kyoto University). *Cx3cr1*^−/−^ mice were generated by crossing *Cx3cr1*^fl/fl^ mice with ZP3-Cre mice (kindly provided by Seiji Hitoshi, Shiga University of Medical Science). CD45.1 congenic mice on a C57BL/6 background and CD45.1/CD45.2 heterozygous mice were bred and maintained in our facility. Male mice aged 6–10 weeks were used throughout. All mice were maintained in a filtered-air laminar-flow enclosure and given standard laboratory food and water. Animal protocols were approved by the Committee of Ethics on Animal Experiment, Shiga University of Medical Science (Approval nos. 2021-11-3, 2023-3-3, 2024-7-6, 2026-2-6).

### Induction of coronary arteritis and treatments

Mice were primed with an intraperitoneal injection of 20 µg of lipopolysaccharide (LPS) on day 0, and injected subcutaneously with 50 µg of FK565 (Peptide Institute) on days 1 and 4, as described previously^8^, with modifications as described here. To deplete NK cells, mice received 100 µl of rabbit anti-asialo GM1 polyclonal antibody (Fujifilm Wako Pure Chemical Corporation, lot WTG4603; 1.2 mg γ-globulin fraction) or the same volume of polyclonal rabbit IgG (Bio X Cell, BE0095, lot 794122M1) on day −1. To inhibit granzyme B, 1 µg of granzyme B inhibitor IV (Sigma-Aldrich) was administered every 24 h from day 0 to day 6. For chemokine neutralization, mice received 250 µg of mouse anti-mouse CCL5 IgG (Bio X Cell) on days 0, 3 and 5 or 50 µg of rabbit anti-CX3CL1 IgG (Torrey Pines Biolabs) on days 0 and 3, alongside respective isotype controls. For calcineurin inhibition, cyclosporine A (Sandimmune; Novartis Pharma) was administered at 5 mg/kg body weight every 12 h from day 0 to day 6. All treatments except FK565 were administered via intraperitoneal injection; the treatment schedule is summarized in Extended Data Fig. 1a.

### Histology and disease scoring

Mice were euthanized and transcardially perfused with phosphate-buffered saline (PBS). Harvested hearts were fixed in 10% formalin, embedded in paraffin and sectioned at a thickness of 4 μm. Coronary arteritis was assessed at a single anatomically defined level: a transverse section through the aortic root. One section per mouse was evaluated, and the value obtained from that section was taken as the value for that animal. Hematoxylin and eosin (H&E)-stained sections were digitized with a NanoZoomer S60V2 slide scanner (Hamamatsu Photonics), and the pericoronary region was measured on the whole-slide images using ImageJ^51^. The inflammation score was calculated as the pericoronary inflammatory area divided by the coronary artery area, taken as the area enclosed by the external elastic lamina of the same vessel; no thresholding or categorical scoring was applied. Measurements were performed independently by two investigators with concordant results.

For quantitative cellular assessment, images were acquired using a BZ-X800 microscope (Keyence). Total cell counts were automatically determined using the Hybrid Cell Count module within the BZ-X Analyzer software (Keyence) and normalized to the corresponding coronary artery area. For immunofluorescence analysis, paraffin-embedded sections were deparaffinized, and antigen retrieval was performed by heating at 95–98 °C in citrate buffer (pH 6.0). After blocking nonspecific binding sites with donkey serum (ImmunoBioScience, IHR8135; 1:100 dilution), sections were stained with rabbit anti-CX3CL1 (Thermo Fisher Scientific, 14-7986-81; 1:100 dilution), followed by Alexa Fluor 555-conjugated anti-rabbit IgG (Thermo Fisher Scientific, A31572; 1:200 dilution) and DAPI. Fluorescence images were acquired on a TCS SP8 confocal microscope (Leica).

### Antibodies and flow cytometry

Anti-mouse CD3ε (145-2C11), anti-mouse CD4 (RM4-5), anti-mouse CD8α (53-6.7), anti-mouse CD16/CD32 (93), anti-mouse B220 (RA3-6B2), anti-mouse NK1.1 (PK136), anti-mouse F4/80 (BM8), anti-mouse Ly6G (1A8), anti-mouse CD11b (M1/70), anti-mouse CD27 (LG.3A10), anti-mouse CD45 (30-F11), anti-mouse CD45.1 (A20), anti-mouse CD45.2 (104), anti-mouse KLRG1 (2F1/KLRG1), anti-mouse T-bet (4B10), anti-mouse CX3CR1 (SA011F11), anti-mouse NFATc1 (7A6), streptavidin-BV421 and Zombie Aqua Fixable viability dye were purchased from BioLegend. Anti-granzyme B (GB11) was from Invitrogen. For flow cytometry, all antibodies were used at a 1:100 dilution, except anti-T-bet, anti-NFATc1 and anti-granzyme B, which were used at 1:50.

Mice were perfused with PBS, and hearts were minced in RPMI-1640 containing 1 mg/ml collagenase II (Worthington). Subsequently, the cell suspensions were shaken at 230 rpm for 1 h at 37 °C and centrifuged at 60 × *g* to separate cardiomyocytes from non-cardiomyocytes. The supernatant containing the non-cardiomyocyte fraction was passed through a 40 μm mesh filter. For peripheral blood, single-cell suspensions were obtained with red-cell lysis, followed by filtration through a 70 μm mesh filter. Single-cell suspensions were incubated with anti-CD16/CD32 for 10 min, followed by staining with monoclonal antibodies for 30 min on ice and washing. For intracellular staining of T-bet and NFATc1, the cells were fixed and permeabilized using the Foxp3/Transcription Factor Staining Buffer Kit (eBioscience). For granzyme B staining, the cells were fixed and permeabilized using a Cytofix/Cytoperm kit (BD Biosciences). Data were acquired on an LSRFortessa (BD Biosciences). Gating strategies are shown in Extended Data Fig. 1c. For imaging flow cytometry, stained samples were analyzed with a FACSDiscover A8 (BD Biosciences). Nuclear translocation of NFATc1 was quantified as the derived correlation parameter between the NFATc1 and nuclear images using the CellView Lens plugin in FlowJo.

### Mixed bone marrow chimeras

Bone marrow (BM) cells were freshly isolated from the tibia and femur of *Cx3cr1*^−/−^ (CD45.2) mice or CD45.1 congenic mice. After erythrocyte lysis, BM cells were counted and mixed at a 1:1 ratio, and 2 × 10^6^ BM cells from each donor genotype were transferred intravenously into irradiated (9 Gy) CD45.1/CD45.2 heterozygous recipient mice. After 7 weeks of reconstitution, the mice were treated with LPS on day 0 and FK565 on days 1 and 4 as described above. On day 7, cardiac cell suspensions were analyzed using flow cytometry to quantify CD45.1 or CD45.2 single-positive NK cells.

### Microarray analysis

Total RNA was isolated from the proximal regions of coronary arteries harvested on day 7 using the miRNeasy Mini Kit (QIAGEN). RNA quality and purity were assessed with a NanoDrop One spectrophotometer and Qubit 2.0 fluorometer (Thermo Fisher Scientific), and RNA integrity with an Agilent 2100 Bioanalyzer (RNA 6000 Nano kit); all samples had RNA integrity numbers of 9.5–9.8. One hundred nanograms of total RNA per sample was amplified and labeled with Cy3. Labeled cRNA was hybridized to a SurePrint G3 Mouse GE microarray (8 × 60K version 2.0; Agilent Technologies) according to the manufacturer’s instructions. All six samples were hybridized to a single slide. Slides were scanned on an Agilent SureScan Microarray Scanner G4900DA and signals extracted with Feature Extraction (version 12.1.1.1). Scanned microarray images were preprocessed using GeneSpring GX software (version 14.9.1; Agilent Technologies); intensities were log2-transformed and normalized by a 75th percentile shift, which centers each array on its own 75th percentile, with no per-gene normalization. Probes flagged as detected in at least two samples of either group were retained (39,001 probes). Normalized expression intensities were summarized at the gene level by averaging multiple probes corresponding to the same gene symbol. Differentially expressed genes (DEGs) between control and FK565-treated mice (n = 3 individual mice per group, each processed separately) were identified using the limma^52^ package (version 3.66.0). *P* values were adjusted for multiple testing using the Benjamini–Hochberg (BH) false discovery rate (FDR) method. Gene Ontology enrichment analysis for Biological Processes was conducted with the clusterProfiler^53^ package (version 4.16.0) on upregulated DEGs (defined as log_2_FC > 1 and FDR < 0.05), using all gene symbols retained after filtering and probe collapsing as the background universe and thresholds of BH-adjusted *P* < 0.05 and *q* < 0.05. Redundant terms were removed with the simplify function (semantic similarity cutoff 0.7, retaining the term with the lowest adjusted *P* value). To estimate immune cell enrichment in the coronary samples, we performed digital deconvolution using the xCell2^54^ package (version 1.0.11). The ImmGenData^55^ reference was used to train the xCell2 object, which was subsequently applied to the gene expression matrix to derive cell-type-specific enrichment scores. *Cx3cl1* expression was compared between groups with the limma moderated *t* test fitting an intensity-dependent trend to the prior variance. The raw and processed data are available in the Gene Expression Omnibus (GEO) under accession code GSE345561.

### Single-cell RNA-sequencing data and references

For single-cell RNA-seq (scRNA-seq) of immune cells from peripheral blood and heart of FK565-treated mice on day 7, 10,000 FACSAria-sorted immune cells (DAPI^−^CD45^+^; pooled cells from 10 individual mice per group in a single experiment were processed separately for blood and heart, yielding one library per tissue per group) were processed using a Chromium Next GEM Single Cell 3′ kit (version 3.1; 10x Genomics) according to the manufacturer’s instructions. Samples were sequenced on the NovaSeq X Plus platform (Illumina). For scRNA-seq of immune cells from peripheral blood and heart of control mice, raw FASTQ files were obtained from GSM6934210^56^ and GSM3488813^57^, respectively. All raw FASTQ files were processed using CellRanger^58^ (version 7.2.0) and aligned to the mm10 reference genome. Aligned sequence data were further analyzed using the Seurat^59^ R package (version 5.1.0). Genes detected in at least three cells and cells with at least 200 detected genes were retained, and cells with more than 10% mitochondrial reads were excluded; no upper thresholds were applied to gene or unique molecular identifier (UMI) counts. Processed data were log-normalized, and doublets were detected and removed using DoubletFinder^60^ (version 2.0.4), and batch effects were corrected using Harmony^61^ (version 1.2.0). Clustering was performed on the first 18 principal components at a resolution of 0.8. NK cells were subclustered on the Harmony embedding using the first 11 principal components at a resolution of 0.5. The raw and processed data are available in the GEO under accession code GSE338096.

For human datasets, data from 11 healthy donors and 13 patients with KD from 5 studies (six healthy donors from GSE166489^26^; three healthy donors and six patients with KD before IVIG from GSE168732^19^; one SARS-CoV-2-negative patient with KD from GSE167029^27^; and two healthy donors and six patients with KD from GSE200743^28^ and GSE254657, unpublished dataset deposited in the Gene Expression Omnibus by the investigators of GSE200743) were integrated. Genes detected in at least three cells and cells with at least 200 detected genes were retained, and upper thresholds for detected genes, UMI counts and mitochondrial content were set individually for each sample on the basis of the distribution of each metric (detected genes, 2,000–6,000; UMI counts, 6,000–20,000; mitochondrial reads, 5–15%; per-sample values are given in Supplementary Table 3). To account for the substantial batch effects and technical heterogeneity across the datasets, each sample was independently normalized with SCTransform^62^, regressing out the percentage of mitochondrial reads. Merged samples were used for computation of principal components with the SCT assay, and batch effects were corrected using Harmony^61^ (version 1.2.0).

Clustering was performed on the first 13 principal components at a resolution of 0.5. In parallel, the RNA assay was log-normalized with a scale factor of 10,000, and the 2,000 most variable features were identified by the vst method followed by scaling of all genes. NK cells were subclustered on the Harmony embedding using the first 13 principal components at a resolution of 0.35. For CX3CL1 expression in human tissue, processed scRNA-seq data were obtained for human coronary arteries^30^ from Zenodo (https://doi.org/10.5281/zenodo.6032099) and for the human vascular cell atlas^31^ from https://www.vascularcellatlas.org/.

DEGs were identified for each NK cell subcluster using the FindAllMarkers function implemented in Seurat on the log-normalized RNA assay with default parameters. Gene Ontology Biological Process enrichment was assessed across clusters with the compareCluster function from the clusterProfiler package, using upregulated DEGs (average log_2_FC > 0.5 and adjusted *P* < 0.05) as input, all genes detected in the dataset as the background universe, and thresholds of BH-adjusted *P* < 0.05 and *q* < 0.05. For the mouse datasets, single-sample gene set enrichment analysis (ssGSEA) for NK subclusters was performed using escape^63^ (version 2.4.0) with the MSigDB mouse hallmark (MH) and Gene Ontology Biological Process (M5) collections together with the BIOCARTA_NFAT_PATHWAY, GOBP_CALCINEURIN_MEDIATED_SIGNALING and NFATC1_TARGET_GENES gene sets.

To infer intercellular communication networks, we applied the CellChat^64^ R package (version 2.2.0) using the CellChatDB.mouse database. Datasets were segregated by experimental condition and tissue origin. Within each subset, communication probabilities for overexpressed ligand–receptor pairs were computed and aggregated into signaling pathways. Network centrality analysis identified predicted dominant cellular sources (outgoing signaling patterns) of key inflammatory and regulatory mediators, and outgoing signaling roles were visualized via ComplexHeatmap^65^ (version 2.25.2). The relative contributions of ligand–receptor pairs were visualized using bar charts, and CCL5–CCR1 communications were illustrated as alluvial plots.

To infer gene regulatory networks for identifying key transcription factors in NK cell subclusters, Seurat-processed data were further analyzed using the SCENIC^66^ R package (version 1.3.1). Following the exclusion of mitochondrial and ribosomal genes, count matrices were filtered to retain genes expressed in >1% of cells. Genes were further filtered with the SCENIC geneFiltering function (minimum total counts, 3% of the number of cells; minimum number of cells detecting the gene, 1%), and gene–gene correlations were computed with runCorrelation to assign regulon directionality. GENIE3^67^ (version 1.30.0) identified co-expression modules from log_2_-transformed data, which RcisTarget (version 1.28.1) refined into regulons using mouse reference databases (mm9-500bp-upstream and mm9-tss-centered-10kb). Only high-confidence regulons (using the top50perTarget method, excluding “extended” targets) were retained. Regulon activity was scored per cell using AUCell (version 1.30.1). Cluster-averaged area under the curve (AUC) scores were *Z*-score normalized and visualized using ComplexHeatmap.

For RNA velocity-based trajectory inference, CellRanger outputs for the mouse datasets were processed using velocyto^68^ (version 0.17.17), and further analyzed using scVelo^69^ (version 0.2.5). RNA velocities were estimated with the stochastic model using default parameters, and velocity vectors were projected onto the UMAP embedding. For Monocle-based trajectory inference, processed Seurat data were analyzed using Monocle3^70,71^ (version 1.3.7). The principal graph was learned without partitioning, with 300 (mouse) or 255 (human) graph nodes and a minimal branch length of 10. Cells were ordered in pseudotime with the root placed in the most immature subcluster (mNK3 in mouse; hNK5 in human).

To assess transcriptional co-regulation while mitigating scRNA-seq sparsity, processed Seurat data were analyzed using hdWGCNA^72^ (version 0.4.12) for metacell-based correlation analysis. For the mouse datasets, metacells were constructed separately within each NK subcluster of each sample, defined by condition and tissue, by aggregating the expression profiles of the 25 nearest neighbors (*k* = 25) in the Harmony embedding, with a maximum of 10 cells shared between metacells. For the human datasets, metacells were constructed separately within each NK subcluster of each donor using the same parameters as for the mouse datasets. For the mNK2 and hNK1 subclusters, pairwise Spearman rank correlation coefficients were calculated between selected transcription factors and effector genes across metacells.

For cross-species integration, human and mouse single-cell datasets were integrated with SATURN^29^ using the 21 coarse cell-type labels (Fig. 4d) and the default parameters, filtering to 8,000 highly variable genes per species and 2,000 macrogenes. The resulting shared embedding was visualized by UMAP using the default parameters in Scanpy^73^ (version 1.10.3). For K-means clustering (*k* = 21, matching the number of coarse cell types), the data were projected onto the leading principal components explaining 90% of the total variance. Differential macrogene expression analysis was performed in Scanpy using the Wilcoxon rank-sum test on the SATURN macrogene values, comparing mNK2 with all other cells in the integrated dataset. Macrogenes were ranked based on their statistical significance, and the top 10 differentially expressed macrogenes were plotted for visualization. For cross-species ortholog mapping and module scoring, mouse–human ortholog relationships were retrieved from Ensembl release 115 using biomaRt^74^ (version 2.66.0). Only pairs annotated as ortholog_one2one with a non-empty human gene symbol were retained, and residual duplicate symbols on either side were removed, yielding 17,092 strictly one-to-one orthologous pairs. This one-to-one ortholog table was used both for the module scores and for the transcriptome-wide correlation. Subcluster signatures were defined with the FindMarkers function (min.pct = 0.25 and logfc.threshold = 0.5) in Seurat on the log-normalized RNA assay, comparing each subcluster with all other subclusters of the same species, and were retained at an adjusted *P* < 0.01. A lower cut-off was applied to human signatures than to mouse signatures (average log_2_ fold change > 0.5 and > 1, respectively) because fold changes were compressed after the integration of five datasets. Signature genes were mapped to the other species through the ortholog table, and genes without a one-to-one ortholog were discarded without replacement. Module scores were calculated with AddModuleScore in Seurat using default parameters. The hNK1 signature was mapped from human to mouse (16 of 25 genes had a one-to-one ortholog) and scored in mouse NK cell subclusters, and the mNK2 and mNK6 signatures were mapped from mouse to human (19 of 31 and 93 of 108 genes, respectively, had a one-to-one ortholog) and scored in human NK subclusters.

### RNA-sequencing data processing and visualization

RNA-seq data of CD27^+^CD11b^+^ cells (stage III) from wild-type and *Tbx21*^−/−^ mice (GSE122874^25^) were processed for quality control and read filtering using fastp^75^ (version 1.0.1). Filtered reads were aligned to the mouse reference genome mm10 using STAR^76^ (version 2.7.10b). Transcriptome-aligned reads were generated using STAR and quantified at the gene level using RSEM^77^ (version 1.3.1) with the estimate-rspd option. Differential expression was analyzed using DESeq2^78^ (version 1.48.2). *P* values were adjusted for multiple testing using the BH method, and genes with an absolute log_2_FC > 1 and an adjusted *P* < 0.05 were considered differentially expressed.

### Epigenomic analysis of the *Tbx21* and *TBX21* loci

To visualize the chromatin landscape of the mouse *Tbx21* locus (mm9, chr11:96,900,000– 97,050,000), the following publicly available datasets were displayed as genome-coverage tracks: NFATc1 bio-ChIP-seq in thymocytes (GSE198031^79^) and cytotoxic CD8^+^ T cells (CTLs; GSE98726^80^); ATAC-seq (GSE77695^37^), STAT4 ChIP-seq (GSE106137^81^), p300 ChIP-seq (GSE77695^37^), and DNase-seq (GSE172358^16^) in NK cells; and c-Jun ChIP-seq in CD4^+^ T and Th17 cells (GSE39756^82^). bigWig tracks mapped to mm10 were lifted over to mm9 using CrossMap^83^ (version 0.7.0); where required, aligned reads in BED format were converted to genome-coverage bedGraph files using bedtools^84^ (version 2.31.1) and then to bigWig using bedGraphToBigWig (University of California, Santa Cruz (UCSC) tools). For human NFATC1 and NFATC3 peak visualization, ChIP-seq datasets for NFATC1 (ENCSR000BQL) and NFATC3 (ENCSR437GBJ) in GM12878 lymphoblastoid cells were used.

To compare the mouse and human loci, human NK cell ATAC-seq data (GSE264693^85^) were processed using fastp for adapter trimming. Reads were aligned to hg19 with bowtie2^86^ (version 2.5.5) and properly paired reads with a mapping quality (MAPQ) ≥ 30 were retained after removal of reads mapping to mitochondrial, random and unplaced contigs; duplicates were removed with samtools (version 1.24) markdup, ENCODE blacklist^87^ regions were removed, and coverage tracks were generated with deepTools (version 3.5.6)^88^ bamCoverage (counts per million (CPM)-normalized, 10-bp bins, with read extension). The mouse *Tbx21 CNS12* region (mm9, chr11:96,988,013–96,988,777) was converted to human coordinates (hg19, chr17:45,798,226–45,799,080) using UCSC liftOver. NFAT, AP-1 and STAT4 motifs were identified with FIMO^89^ (version 5.5.9) using JASPAR^90^ position-weight matrices for NFATC1 (MA0624.1), NFATC2 (MA0152.1), NFATC3 (MA0625.2), FOS::JUN(MA0099.3), and STAT4 (MA0518.1) under the default background model (*P* < 1 × 10^−4^), and at a relaxed threshold of *P* < 1 × 10^−3^. In each species, the matrices were scanned across the CNS12 region extended by 500 base pairs on each side, and occurrences falling within the orthologous *CNS12* were reported. All motifs except FOS::JUN (AP-1) were detected at *P* < 1 × 10^−4^; the FOS::JUN (AP-1) motif was detected in both species only at the relaxed threshold. Integrated multiomics tracks across the *Tbx21* and *TBX21* loci were visualized using IGV^91^ (version 2.19.7).

### Statistical analysis

Measurements analyzed in GraphPad Prism (version 8) are summarized as mean ± s.e.m. Data distribution was assumed to be normal, but this was not formally tested. Two-tailed unpaired *t* tests were used for comparisons between two independent groups. Two-tailed paired or ratio paired *t* tests were used for comparisons between two paired groups; ratio paired tests were applied to fluorescence intensity measurements. One-way analysis of variance (ANOVA) followed by a Tukey or Dunnett post hoc multiple comparisons test was used for comparisons of multiple groups. Two-way ANOVA with repeated measures on the within-animal factor and a Geisser–Greenhouse correction, followed by a Šídák post hoc multiple comparisons test, was used for designs combining a within-animal factor with a between-animal factor. For scRNA-seq data, DEGs and subcluster signatures were identified with the two-sided Wilcoxon rank-sum test implemented in Seurat, with Bonferroni-adjusted *P* values. Differentially expressed macrogenes were identified with the Wilcoxon rank-sum test in Scanpy. Differences were considered significant at *P* < 0.05. Exact *P* values are shown in the figures; t statistics with degrees of freedom and, for multiple-comparison tests, mean differences with 95% confidence intervals are provided in the Source Data.

## Reporting summary

Further information on research design is available in the Nature Portfolio Reporting Summary linked to this article.

## Data availability

The single-cell RNA-seq and microarray data generated in this study have been deposited in the Gene Expression Omnibus (GEO) under accession codes GSE338096 and GSE345561. Publicly available datasets analyzed in this study are available in the GEO under accession codes GSE166489, GSE168732, GSE167029, GSE200743, GSE254657, GSE122874, GSE222854 (GSM6934210), GSE122930 (GSM3488813), GSE77695, GSE106137, GSE172358, GSE198031, GSE98726, GSE39756 and GSE264693, and in ENCODE under accession codes ENCSR000BQL and ENCSR437GBJ. Additional processed datasets are available at Zenodo (https://doi.org/10.5281/zenodo.6032099) and at https://www.vascularcellatlas.org/. Source data are provided with this paper.

## Code availability

Analyses used publicly available software as described in Methods; no custom code was used.

## Acknowledgements

This work was supported by JSPS (Japan Society for the Promotion of Science) KAKENHI Grant Numbers JP22K20757 (M.F.), JP23K07224 (S.H.), JP23K06498 (H.S.), JP26K09989 (H.S.), JP22K07132 (T.H.), JP25K10411 (T.H.), and JP22H04922 (AdAMS; M.F.), by The Mother and Child Health Foundation (H.S.), and by Kawano Masanori Memorial Public Interest Incorporated Foundation for Promotion of Pediatrics (H.S.). We thank Robin James Storer, PhD, from Edanz for editing and reviewing drafts of this manuscript.

## Author information

These authors contributed equally: Masami Fujita, Hiroki Satooka.

## Contributions

M.F., H.S. and T.H. conceived and designed the study. M.F. and H.S. performed experiments and analyzed results. S.H. prepared the samples for the microarray assays. H.S., S.M. and E.J.V. performed the bioinformatics analyses. M.F., H.S., Y.M. and T.H. discussed the results of experiments. K.K., Y.A. and M.E. provided key resources. M.F., H.S. and T.H. wrote the manuscript, and all authors read, revised and approved the final version, and agree both to be personally accountable for their own contributions and to ensure that questions related to the accuracy or integrity of any part of the work, even ones in which they were not personally involved, are appropriately investigated, resolved and the resolution documented in the literature.

## Corresponding authors

Correspondence to Hiroki Satooka or Takako Hirata.

