## Extended Data Figures for "Terminally differentiated CX3CR1^+^ natural killer cells drive coronary arteritis through an NFAT–T-bet axis"

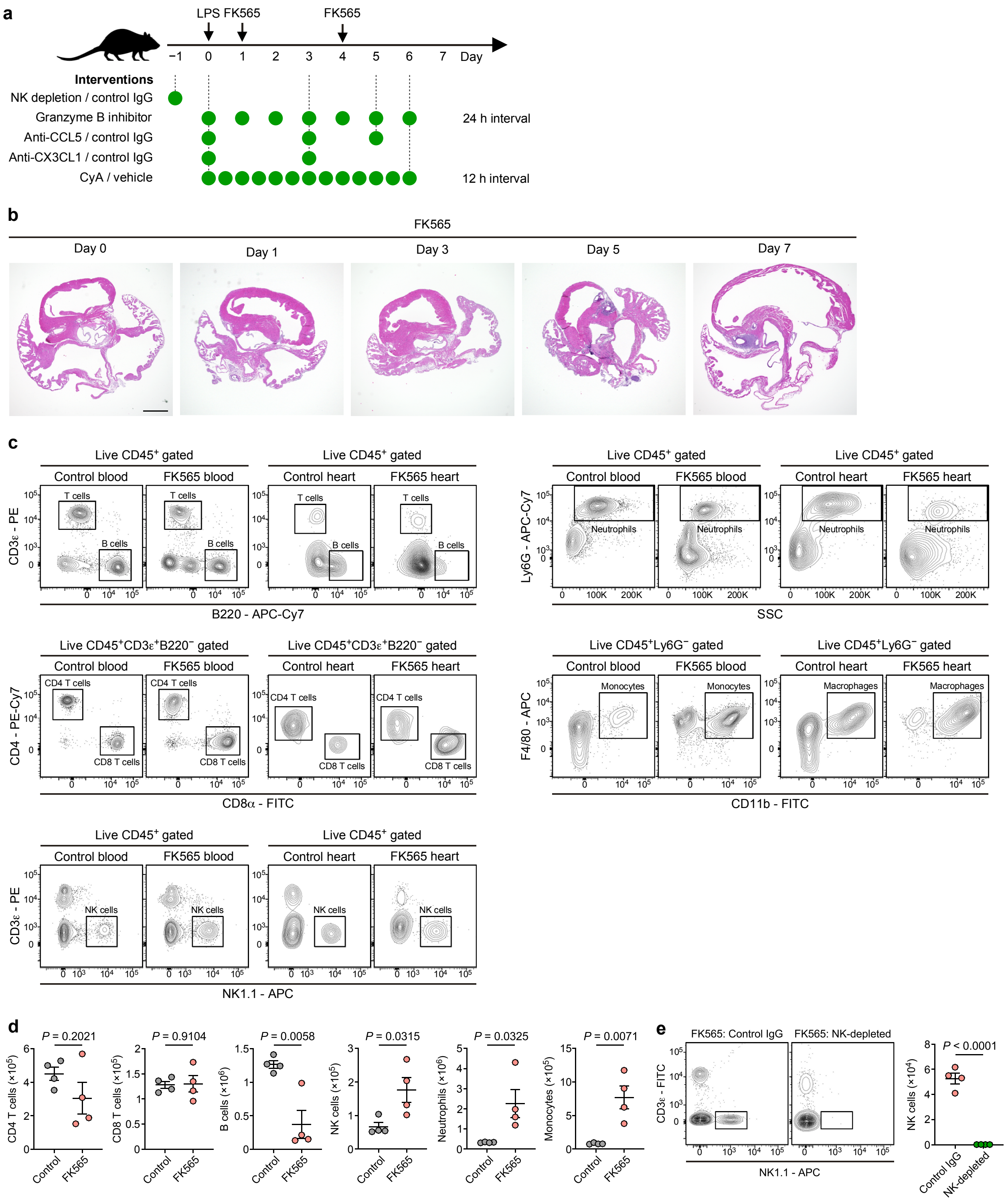


**Extended Data Fig. 1 | Induction of coronary arteritis and systemic immune cell changes in FK565-treated mice. a**, Schematic of the experimental timeline described in Methods. **b**, Representative images of H&E staining of the coronary arteries and the aortic roots from FK565-treated mice on days 0, 1, 3, 5 and 7. Scale bar, 500 μm. **c**, Gating strategy for flow cytometric analysis of peripheral blood and heart from control or FK565-treated mice. **d**, Flow cytometric quantification of immune cell populations in the peripheral blood of control and FK565-treated mice. Cell types were identified as follows: CD4 T cells ($CD45^+CD3^+CD4^+CD8^−$), CD8 T cells ($CD45^+CD3^+CD4^−CD8^+$), B cells ($CD45^+CD3^−B220^+$), NK cells ($CD45^+CD3^−NK1.1^+$), neutrophils ($CD45^+Ly6G^+$), monocytes ($CD45^+Ly6G^−F4/80^+$) ($n = 4$). **e**, Flow cytometry plot (left) and quantification (right) of NK cells in peripheral blood of FK565-treated mice given control IgG or NK-depleting antibody on day 7 ($n = 4$). Data are mean ± s.e.m., and individual dots represent individual mice. Statistical significance was determined using two-tailed unpaired *t* tests for **d** and **e**. Data are representative of more than three independent experiments.

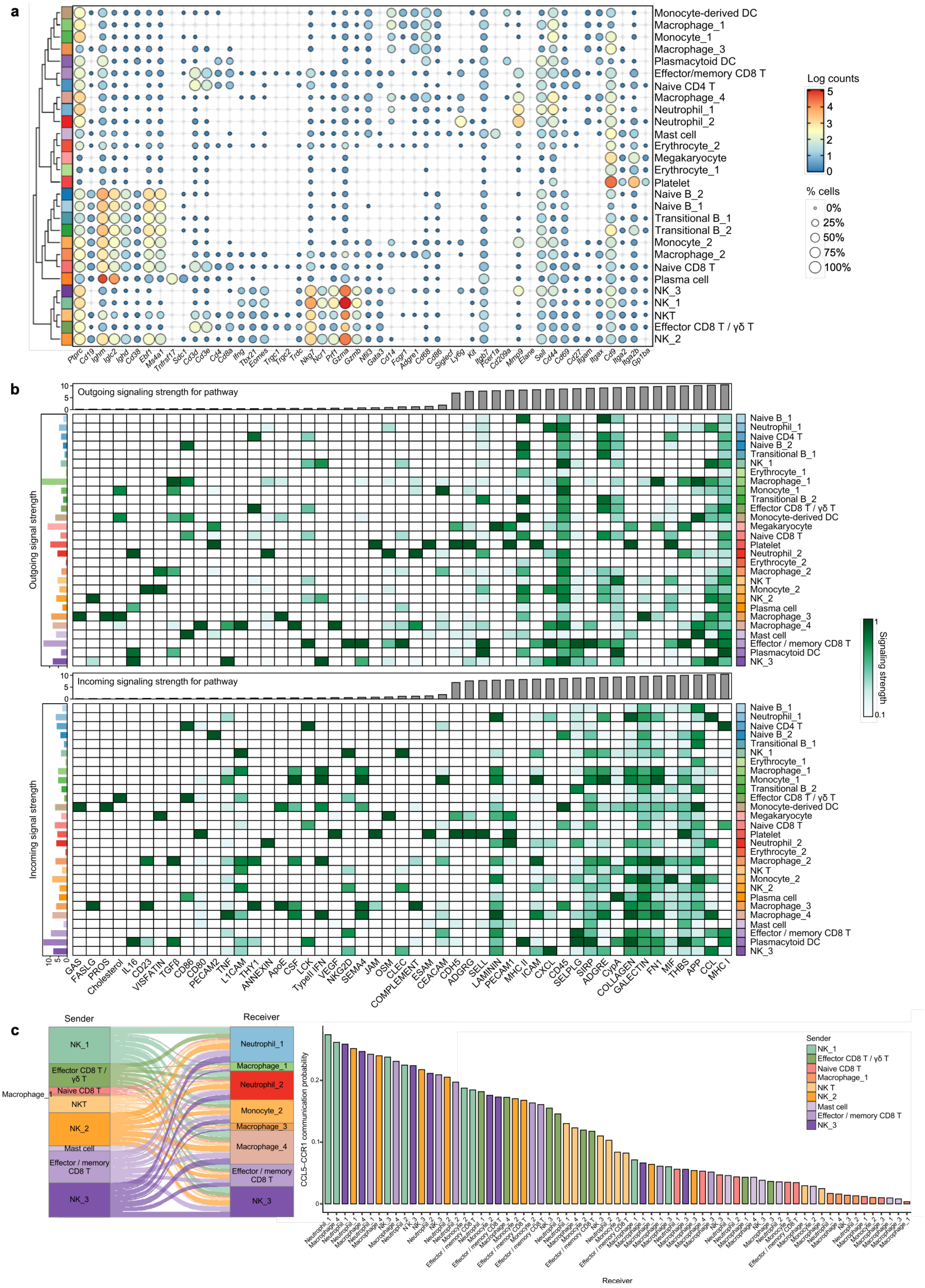
a
Monocyte-derived DC
Macrophage_1
Monocyte_1
Macrophage_3
Plasmacytoid DC
Effector/memory CD8 T
Naive CD4 T
Macrophage_4
Neutrophil_1
Neutrophil_2
Mast cell
Erythrocyte_2
Megakaryocyte
Erythrocyte_1
Platelet
Naive B_2
Naive B_1
Transitional B_1
Transitional B_2
Monocyte_2
Macrophage_2
Naive CD8 T
Plasma cell
NK_3
NK_1
NKT
Effector CD8 T / γδ T
NK_2
Log counts
5
4
3
2
1
0
% cells
0%
25%
50%
75%
100%
Ptprc Cd19 Ighm Iglc2 Ighd Cd38 Ebf1 Ms4a1 Tnfrsf17 Sdc1 Cd3d Cd3e Cd4 Cd8a Ifng Tbx21 Eomes Trgc1 Trgc2 Trdc Nkg7 Ncr1 Prf1 Gzma Gzmb Nfil3 Gata3 Cd14 Fcgr1 Adgre1 Cd68 Cd86 Siglecf Ly6g Kit Itgb7 Fcer1a Cd209a Mmp9 Elane Sell Cd44 Cd69 Cd27 Itgam Itgax Cd9 Itga2 Itga2b Gp1ba
b
Outgoing signaling strength for pathway
Outgoing signal strength
Incoming signaling strength for pathway
Incoming signal strength
Signaling strength
1
0.1
Naive B_1
Neutrophil_1
Naive CD4 T
Naive B_2
Transitional B_1
NK_1
Erythrocyte_1
Macrophage_1
Monocyte_1
Transitional B_2
Effector CD8 T / γδ T
Monocyte-derived DC
Megakaryocyte
Naive CD8 T
Platelet
Neutrophil_2
Erythrocyte_2
Macrophage_2
NK T
Monocyte_2
NK_2
Plasma cell
Macrophage_3
Macrophage_4
Mast cell
Effector / memory CD8 T
Plasmacytoid DC
NK_3
GAS FASLG PROS Cholesterol IL16 CD23 VISFATIN TGFβ CD86 CD80 PECAM2 TNF L1CAM THY1 ANNEXIN ApoE CSF LCK TypeII IFN VEGF NKG2D SEMA4 JAM OSM CLEC COMPLEMENT ESAM CEACAM CDH5 ADGRG SELL LAMININ PECAM1 MHC II ICAM CXCL CD45 SELPLG SIRP ADGRE CypA COLLAGEN GALECTIN FN1 MIF THBS APP CCL MHC I
c
Sender
Receiver
NK_1
Effector CD8 T / γδ T
Naive CD8 T
Macrophage_1
NKT
NK_2
Mast cell
Effector / memory CD8 T
NK_3
Neutrophil_1
Macrophage_1
Neutrophil_2
Monocyte_2
Macrophage_3
Macrophage_4
Effector / memory CD8 T
NK_3
CCL5–CCR1 communication probability
Sender
NK_1
Effector CD8 T / γδ T
Naive CD8 T
Macrophage_1
NK T
NK_2
Mast cell
Effector / memory CD8 T
NK_3
Receiver

**Extended Data Fig. 2 | Intercellular communication analysis reveals NK cells as dominant senders of CCL5–CCR1 signaling in FK565-induced coronary arteritis. a**, Dot plot of selected marker genes across annotated immune cell clusters from peripheral blood and heart of control or FK565-treated mice. **b**, Heatmaps showing outgoing (top) and incoming (bottom) signaling strength for each signaling pathway across cell populations identified in the heart of FK565-treated mice, computed by CellChat network centrality analysis (out-degree and in-degree, respectively). Color intensity is scaled within each pathway. Bar plots above each heatmap indicate the total signaling strength summed across all pathways for each cell population; bar plots to the right indicate the total strength summed across all cell populations for each pathway. **c**, Alluvial plot showing inferred CCL5–CCR1 signaling from sender to receiver populations. Ribbon width represents the summed communication probability (left). Communication probability for each sender–receiver pair, ranked in descending order. Bars are colored by sender population (right).

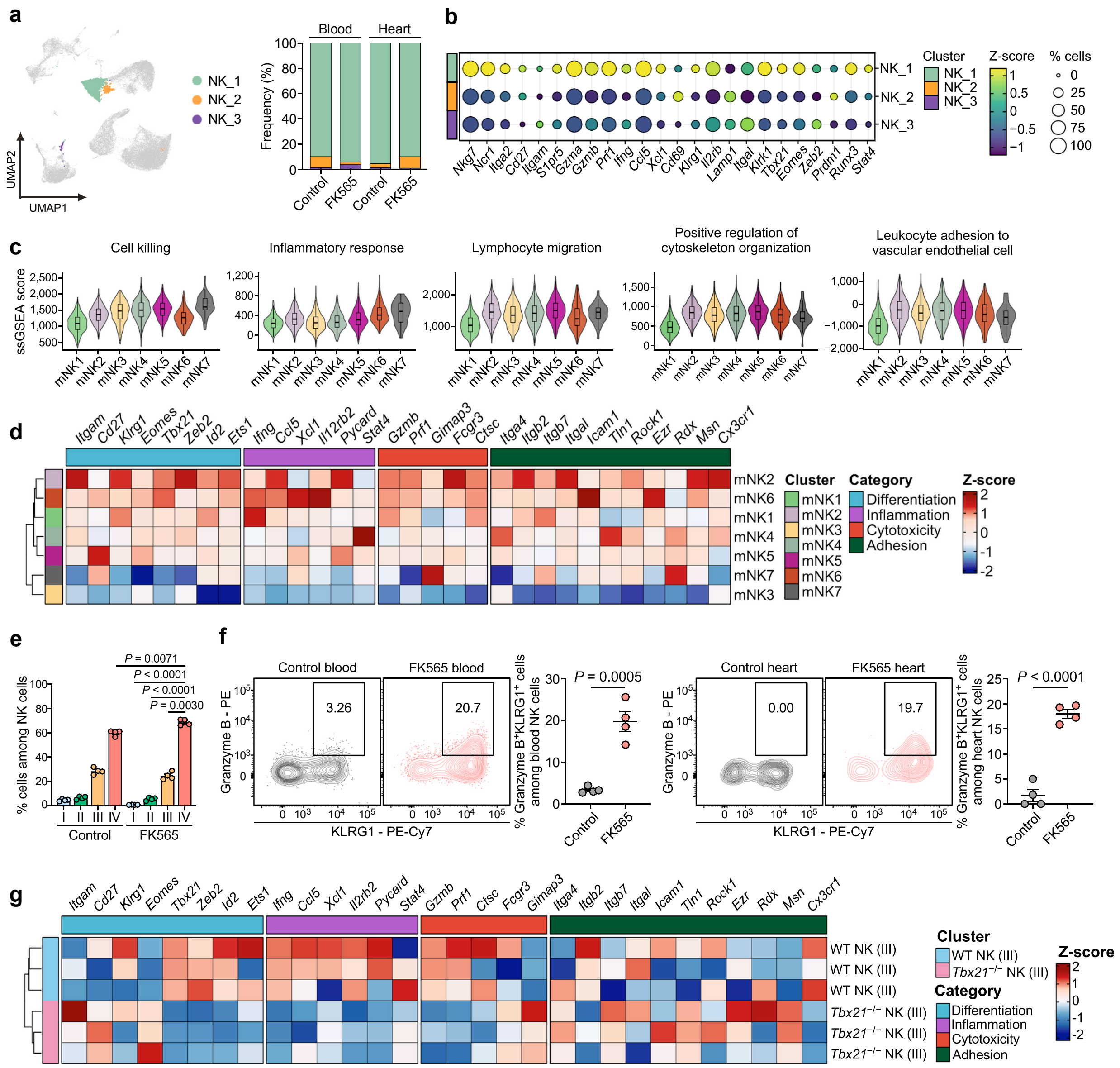


**Extended Data Fig. 3 | Functional classification of NK cell subsets revealed enhanced cytotoxicity and migratory capacity in FK565-treated mice. a**, NK cell clusters were highlighted in UMAP embedding of immune cells from peripheral blood and heart of control or FK565-treated mice. **b**, Dot plot showing the expression of selected NK cell-related genes across NK cell clusters. **c**, Violin plots illustrating the distribution of ssGSEA scores for the indicated functional modules across mNK cell subclusters from peripheral blood and heart of control or FK565-treated mice. **d**, Pseudobulk heatmap depicting the expression of selected genes associated with the indicated biological processes across NK cell subclusters. **e**, Frequencies of NK cell maturation stages (stage I, $CD27^{-}CD11b^{-}$; stage II, $CD27^{+}CD11b^{-}$; stage III, $CD27^{+}CD11b^{+}$; stage

IV, CD27$^{-}$CD11b$^{+}$) among blood NK cells from control and FK565-treated mice on day 7, determined by flow cytometry ($n$ = 4 per group). **f**, Representative flow cytometry plots and quantification of granzyme B$^{+}$KLRG1$^{+}$ cells among NK cells in the blood and heart of control and FK565-treated mice on day 7 ($n$ = 4 per group). Numbers in plots indicate the percentage of cells in the gate. **g**, Heatmap depicting the expression of selected genes associated with the indicated biological processes across mouse NK cell maturation stage III (CD27$^{+}$CD11b$^{+}$) from wild-type (WT) and T-bet-deficient (*Tbx21*$^{-/-}$) mice (GSE122874). Data are mean ± s.e.m., and individual dots represent individual mice. Statistical significance in **e** and **f** was determined using a two-way ANOVA with repeated measures on maturation stage and the Geisser–Greenhouse correction followed by Šídák multiple comparisons test for **e** and two-tailed unpaired $t$ tests for **f**. Data are representative of more than three independent experiments.

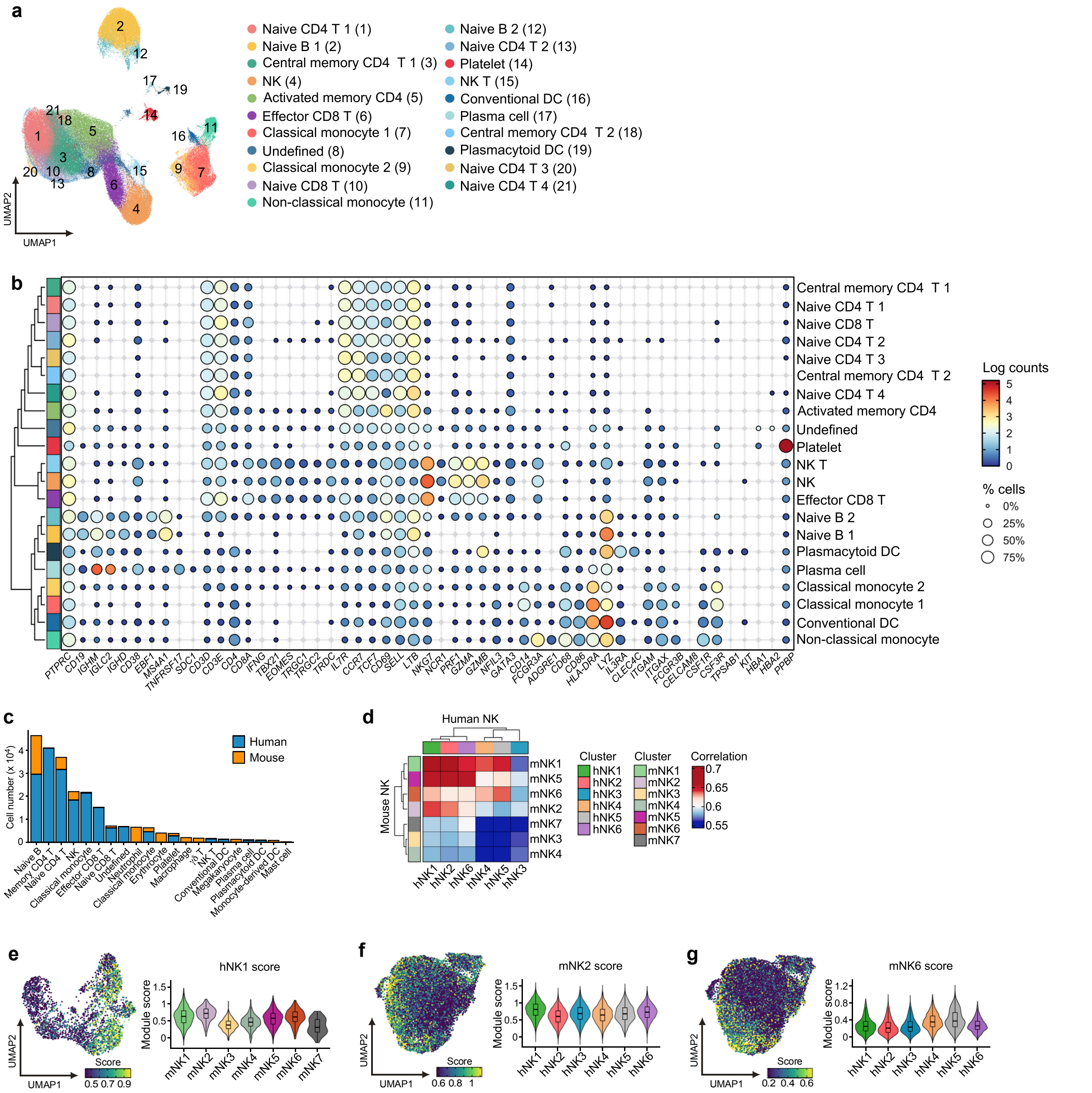


**Extended Data Fig. 4 | Single-cell RNA-seq profiling of immune cell populations and cross-species comparison of human and mouse NK cells. a** and **b**, UMAP (a) embedding of immune cells (181,728 cells across 21 cell types) and dot plot (b) of selected marker genes across annotated immune cell clusters from peripheral blood of healthy controls and patients with Kawasaki disease. **c**, Number of cells of each coarse cell type per species in the SATURN integration. **d**, Heatmap showing the pairwise Spearman correlation of global gene expression profiles between the identified mouse (mNK1–mNK7) and human (hNK1–hNK6) NK cell subclusters. To enable direct cross-species comparison, mouse genes were mapped to their unique one-to-one human orthologs using the Ensembl biomaRt database. Pseudobulk average expression profiles were generated for each subcluster, and genes with a mean expression > 0.5 and a variance > 0.05 were retained for correlation analysis. **e**, Feature plot displaying the hNK1 signature module score projected onto the UMAP embedding of the mNK subclusters, identified in **Fig. 3a** (left). hNK1 signature genes (average $log_2FC > 0.5$, adjusted $P < 0.01$ compared with all other human NK subclusters) were mapped from human to mouse using the one-to-one ortholog table described in Methods. Violin plots illustrating the distribution of the signature module scores across the defined mNK cell subclusters (right). **f**, Feature plot displaying the mNK2 signature module score projected onto the UMAP embedding of the hNK subclusters (left). mNK2 signature genes (average $log_2FC > 1$, adjusted $P < 0.01$ compared with all other mouse NK subclusters) were mapped from mouse to human using the one-to-one ortholog table described in Methods. Violin plots illustrating the distribution of the signature module scores across the defined hNK cell subclusters (right). **g**, Feature plot displaying the mNK6 signature module score projected onto the UMAP embedding of the hNK subclusters (left). mNK6 signature genes (average $log_2FC > 1$, adjusted $P < 0.01$ compared with all other mouse NK subclusters) were mapped from mouse to human using the one-to-one ortholog table described in Methods. Violin plots illustrating the distribution of the signature module scores across the defined hNK cell subclusters (right). To minimize the visual impact of extreme outliers, the color scale limits were set to the 30th and 95th percentiles.

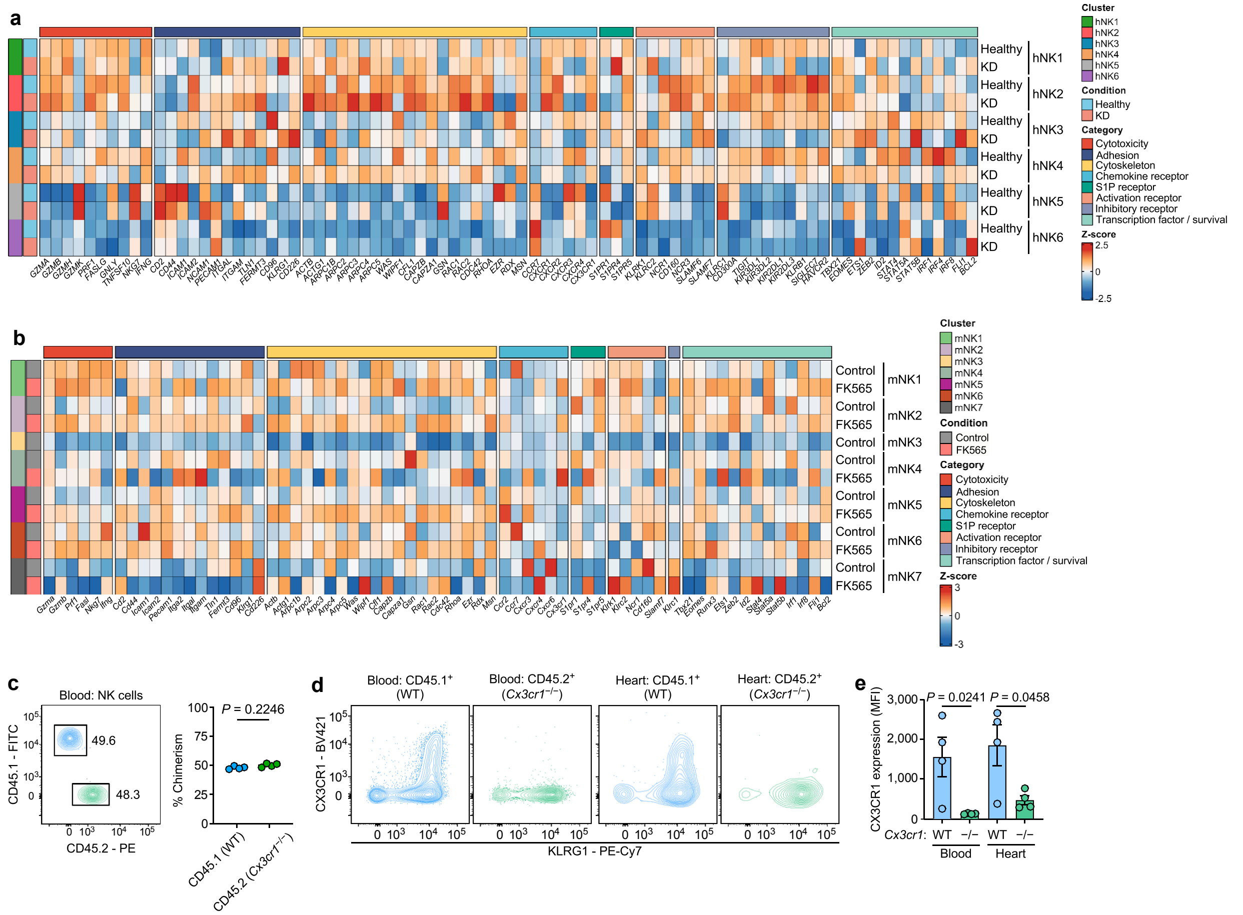


**Extended Data Fig. 5 | Shared transcriptional signatures of NK cell subsets in human Kawasaki disease and the FK565-induced mouse model. a** and **b**, Heatmap of the expression of selected genes associated with the indicated biological processes across human NK cell subclusters (hNK1–hNK6) in healthy controls and patients with Kawasaki disease (a) and of mouse NK cell subclusters (mNK1–mNK7) in control and FK565-treated mice (b). Expression values were averaged within each subcluster and condition, and *Z*-score normalized for each gene. **c**, Representative flow cytometry plot (left) and quantification (right) of the proportions of wild-type (CD45.1$^{+}$) and *Cx3cr1*$^{-/-}$ (CD45.2$^{+}$) NK cells in the blood of mixed bone marrow chimeras on day 0, showing approximately equal donor contributions at baseline. Numbers in the plot indicate the percentage of cells in each gate. **d**, Representative flow cytometry plots of CX3CR1 and KLRG1 expression in wild-type (CD45.1$^{+}$) and *Cx3cr1*$^{-/-}$ (CD45.2$^{+}$) NK cells from the blood and heart of FK565-treated mixed bone marrow chimeras on day 7. **e**, Quantification of CX3CR1 expression in the populations shown in **d**, confirming loss of CX3CR1 in *Cx3cr1*$^{-/-}$ cells. Data in **c** and **e** are mean ± s.e.m., and individual dots represent individual mice (*n* = 4). Statistical significance was determined by two-tailed paired *t* tests for **c** and two-tailed ratio paired *t* tests for **e**, reflecting the measurement of both donor populations within each chimeric mouse. Data are representative of three independent experiments.

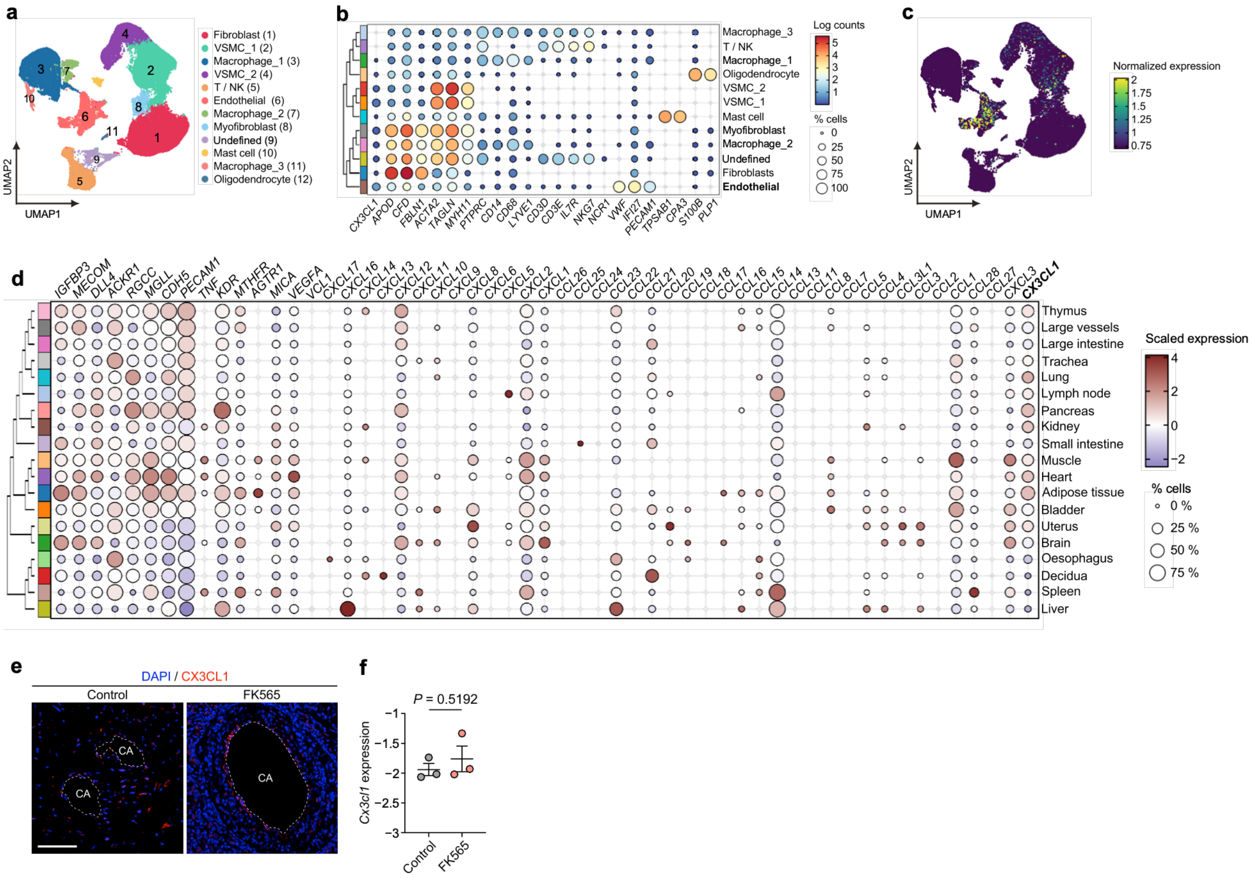


**Extended Data Fig. 6 | CX3CL1 expression marks the coronary endothelium as a target for CX3CR1⁺ NK cells.** **a**, UMAP embedding of cells from human coronary artery (23,148 cells across 12 cell types). **b**, Dot plot of selected marker genes across annotated cell clusters. **c**, Expression of *CX3CL1* in annotated clusters. **d**, Dot plot of chemokine and endothelial marker gene expression across endothelial cells from the indicated human tissues reported as human vascular endothelial cell atlas. **e**, Representative immunofluorescence images (left) of coronary arteries from control and FK565-treated mice on day 7, stained for CX3CL1 (red) and DAPI (blue). Dashed line outlines the vessel lumen. Scale bar, 50 µm. CA = coronary artery. **f**, Quantification of *Cx3cl1* expression in coronary arteries from control and FK565-treated mice, determined by microarray analysis. Values are $\log_2$(signal / 75th percentile of each array) after 75th percentile shift normalization; negative values reflect expression below the array's 75th percentile. Data in **f** are presented as mean ± s.e.m., and individual dots represent individual mice ($n = 3$). Statistical significance was determined using the moderated $t$ test implemented in limma; BH-adjusted $P$ values are reported.

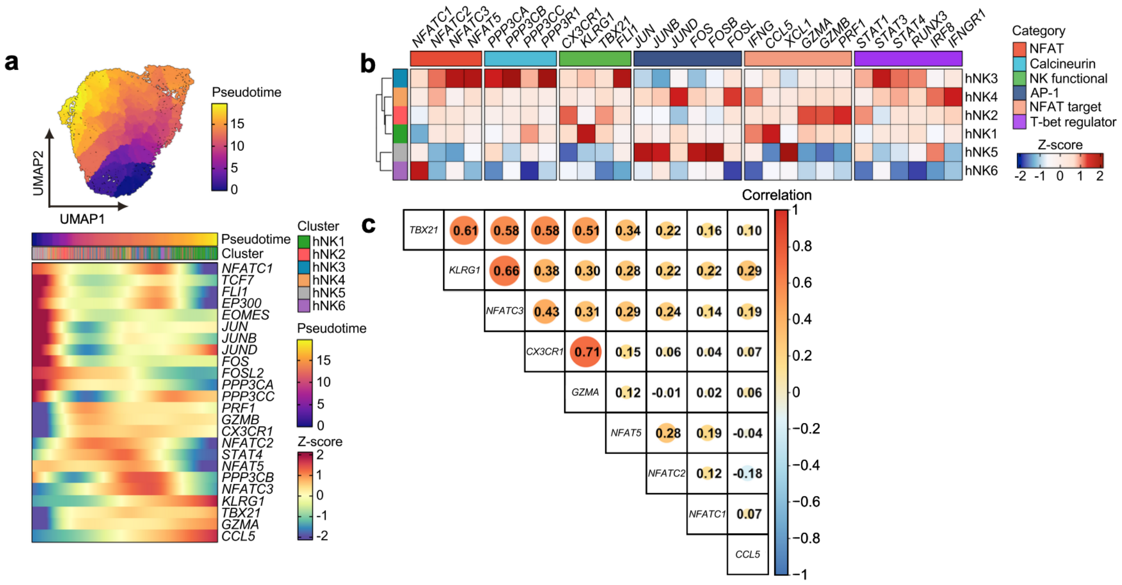

**Extended Data Fig. 7 | NFAT expression accompanies the T-bet effector program in NK cells from patients with Kawasaki disease. a**, Trajectory inference of identified hNK subclusters identified from **Fig. 4a**. hNK subclusters from peripheral blood of patients with Kawasaki disease were analyzed using pseudotime inference with Monocle3. UMAP projections overlaid with pseudotime scores, revealing the dynamic transitions of NK cell subclusters (top). The expression dynamics of NK cell- and NFAT-related genes along the inferred trajectories (bottom). **b**, Heatmap visualizing the pseudobulk gene expression of selected markers for the calcineurin–NFAT pathway, AP-1 transcription factor complex and NK cell functional regulators across hNK cell subclusters (hNK1–hNK6). Gene expression levels were aggregated per subcluster and subsequently scaled per gene to compute Z-scores. **c**, Pairwise Spearman correlation coefficients between the indicated genes across hNK1 metacells, constructed using hdWGCNA (k = 25 nearest neighbors), aggregated separately for each donor. Circle size and color indicate the correlation coefficient.

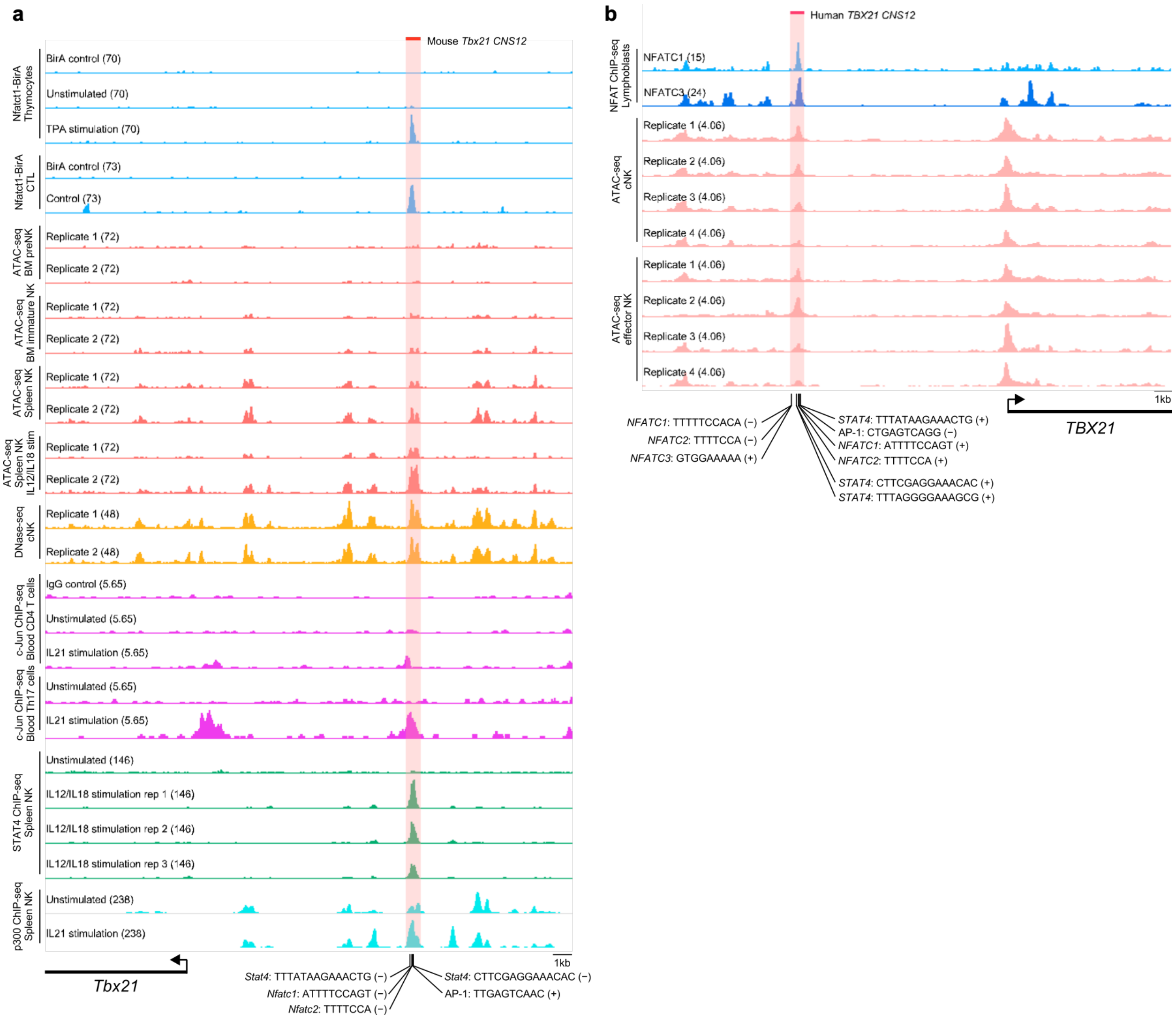


**Extended Data Fig. 8 | An NFAT motif in the *Tbx21*/*TBX21* locus in a region that becomes accessible upon NK cell activation in both mouse and human. a**, An NFAT-binding motif upstream of *Tbx21* locus (mm9) is located in a region that becomes accessible upon NK cell activation. From top: NFATc1-BirA bio-ChIP-seq in thymocytes and in cytotoxic CD8 T cells (CTL); ATAC-seq in NK cells (BM, bone marrow; preNK, premature NK cell; two replicates each); DNase-seq in conventional NK cells (cNK); c-Jun ChIP-seq in CD4 T and Th17 cells; and STAT4 and p300 ChIP-seq in NK cells. All datasets are publicly available (see Methods) and were lifted over to mm9 where required. Positions of NFAT-, STAT4- and AP-1-binding motifs identified within the locus are indicated below, with the corresponding motif sequences shown. **b**, An NFAT-binding motif upstream of the *TBX21* locus (hg19) is located in a region that becomes accessible upon NK cell activation. NFATC1 and NFATC3 ChIP-seq in GM12878 lymphoblastoid cells and ATAC-seq in human NK cell subsets are shown. Positions of NFAT-, STAT4- and AP-1-binding motifs identified within the locus are indicated below, with the corresponding motif sequences shown. The shaded regions correspond to mouse *Tbx21 CNS12* (**a**) and human *TBX21 CNS12* (**b**), the latter converted from mouse coordinates using liftOver (mm9 to hg19). Motifs were identified with FIMO at $P < 1 \times 10^{-4}$, except the AP-1 motif, which was detected at $P < 1 \times 10^{-3}$ and lies 47 bp (mouse) and 49 bp (human) from the NFAT site. The data range is shown in brackets for each track.
