## Supplementary Table 1 for "Terminally differentiated CX3CR1^+^ natural killer cells drive coronary arteritis through an NFAT–T-bet axis"

**Supplementary table 1. SATURN mNK2 macrogenes**

| macrogene 854 | | macrogene 107 | | macrogene 425 | | macrogene 773 | | macrogene 236 | | macrogene 55 | | macrogene 1930 | | macrogene 1410 | | macrogene 1141 | | macrogene 609 | |
|---|---|---|---|---|---|---|---|---|---|---|---|---|---|---|---|---|---|---|---|
| gene | weight | gene | weight | gene | weight | gene | weight | gene | weight | gene | weight | gene | weight | gene | weight | gene | weight | gene | weight |
| mouse_Klra6 | 1.103136 | mouse_Cd69 | 2.025012 | mouse_Tnfrsf9 | 0.267511 | human_GPRC5A | 0.584649 | human_PRSS2 | 0.681125 | human_CFD | 1.149127 | human_GPHB5 | 0.381845 | mouse_Pi16 | 0.189954 | mouse_Nkg7 | 1.270247 | human_SLC22A9 | 0.510017 |
| mouse_Klra5 | 0.901063 | human_CD69 | 1.100722 | mouse_Crim1 | 0.244427 | human_GPRC5D | 0.579293 | human_CELA1 | 0.667593 | human_GZMA | 0.653385 | mouse_Ccl4 | 0.160476 | mouse_Gzma | 0.126483 | human_NKG7 | 0.839877 | human_SLC22A25 | 0.439868 |
| mouse_Klra1 | 0.885991 | human_CLEC1B | 1.07759 | mouse_Xcl1 | 0.224484 | mouse_Nkg7 | 0.532337 | mouse_Cela1 | 0.551103 | human_GZMM | 0.562736 | human_CCL3 | 0.086516 | mouse_Glipr1 | 0.124838 | mouse_Cldnd2 | 0.581803 | human_SLC22A6 | 0.420933 |
| mouse_Klra7 | 0.812471 | mouse_Clec2i | 0.959387 | mouse_Igfbp4 | 0.146025 | mouse_Ms4a6b | 0.39176 | human_CELA2A | 0.48114 | human_PRSS57 | 0.470614 | mouse_Ccl3 | 0.053817 | mouse_Gzmb | 0.124755 | mouse_Tmem204 | 0.456793 | human_SLC22A24 | 0.353899 |
| mouse_Klra3 | 0.732097 | mouse_Klrk1 | 0.676631 | human_XCL1 | 0.120101 | mouse_Rhd | 0.279169 | mouse_Gzmb | 0.367227 | mouse_Gzma | 0.402537 | mouse_Ccl5 | 0.044304 | mouse_Ccl5 | 0.122483 | human_EMP3 | 0.263636 | human_HTR3E | 0.213434 |
| human_KLRB1 | 0.722147 | mouse_Klrb1c | 0.559162 | human_TNFRSF9 | 0.115353 | mouse_Tmem80 | 0.245598 | human_PRSS3 | 0.344362 | human_PRSS33 | 0.333467 | mouse_Serpinb6b | 0.024785 | mouse_Cma1 | 0.100804 | mouse_Tmem37 | 0.210862 | mouse_Ccl5 | 0.211795 |
| mouse_Klra9 | 0.719886 | human_CLEC2A | 0.48554 | human_CHRDL1 | 0.104226 | mouse_Gprc5d | 0.151989 | mouse_Gzma | 0.204238 | human_PRSS41 | 0.324023 | mouse_Cst7 | 0.023823 | mouse_Serpinb9 | 0.096214 | mouse_Emp3 | 0.206085 | human_MOGAT3 | 0.114266 |
| mouse_Klra8 | 0.687445 | mouse_Klrd1 | 0.428768 | human_RSPO3 | 0.089771 | mouse_Gpr180 | 0.120327 | mouse_Cma1 | 0.196393 | human_TPSD1 | 0.319113 | human_SERPINB9 | 0.015609 | mouse_Glipr2 | 0.073723 | human_CMTM6 | 0.167925 | human_CYP2F1 | 0.095332 |
| mouse_Klra4 | 0.678423 | mouse_Klra8 | 0.390359 | human_IL23R | 0.086212 | human_MAL2 | 0.0581 | human_GZMK | 0.191395 | human_TPSAB1 | 0.254252 | human_CCL3L3 | 0.014603 | human_SERPINA1 | 0.064604 | human_TMEM204 | 0.132621 | human_NT5DC4 | 0.086433 |
| human_KLRC1 | 0.615204 | mouse_Klrb1a | 0.388085 | human_CRIM1 | 0.070635 | mouse_Tmem60 | 0.057162 | human_PGA3 | 0.187569 | mouse_Cfd | 0.245581 | mouse_S100a10 | 0.007761 | mouse_Crisp2 | 0.062706 | human_TMEM127 | 0.103067 | mouse_Slc22a12 | 0.0861 |
