## Supplementary Table 2 for "Terminally differentiated CX3CR1^+^ natural killer cells drive coronary arteritis through an NFAT–T-bet axis"

**Supplementary table 2. SATURN mNK2 macrogene expression**

| | mNK2 macrogenes | | | | | | | | | |
|---|---|---|---|---|---|---|---|---|---|---|
| | **609** | **1141** | **1410** | **1930** | **55** | **236** | **773** | **425** | **107** | **854** |
| **Classical monocyte** | 4.92E-05 | 0.13189732 | 0.000323746 | 0.00171823 | 0.26858094 | 0.000378365 | 0.000363608 | 9.06E-06 | 0.044474818 | 0.111390546 |
| **Conventional DC** | 0 | 0.11318699 | 2.07E-05 | 0 | 0.082849935 | 0 | 0 | 0 | 0.09114131 | 0.04794301 |
| **Effector CD8 T** | 0.0003504 | 1.1625088 | 0.001113911 | 8.70E-05 | 0.48687968 | 0.011209291 | 0.001917399 | 0.000375124 | 0.5067509 | 0.37672135 |
| **Erythrocyte** | 0.038960002 | 0.16683869 | 0.07525202 | 0.12019885 | 0.15119398 | 0.086750776 | 0.10928479 | 0.048598945 | 0.0876347 | 0.24103166 |
| **γδ T** | 0.5087802 | 1.7572235 | 0.38727275 | 0.10880438 | 0.33810392 | 0.23849693 | 0.93781525 | 0.3032379 | 1.0974383 | 1.1516767 |
| **Macrophage** | 0.00121973 | 0.06715226 | 0.011692872 | 0.01606037 | 0.008754935 | 0.004325127 | 0.112652525 | 0.000617331 | 0.07610558 | 0.03887735 |
| **Mast cell** | 0.04002072 | 0.21624383 | 0.042633843 | 0.13150945 | 0.03773331 | 0.029648002 | 0.15489988 | 0.012000197 | 0.4585115 | 0.1186447 |
| **Megakaryocyte** | 0.006663218 | 0.073966615 | 0.025726547 | 0.063007355 | 0.08623054 | 0.10425343 | 0.048103347 | 0.019619899 | 0.03995436 | 0.17059027 |
| **Memory CD4 T** | 0 | 0.091886 | 4.69E-06 | 1.54E-06 | 0.09751033 | 0.005674161 | 1.14E-05 | 9.73E-06 | 0.47527492 | 0.10060681 |
| **Monocyte-derived DC** | 0.007435068 | 0.06237841 | 0.008749826 | 0.05350915 | 0.01110201 | 0.007552759 | 0.08056649 | 0.003357875 | 0.19167835 | 0.06695322 |
| **NK T** | 0.020826912 | 0.91998756 | 0.047187965 | 0.014472869 | 0.49706784 | 0.055079915 | 0.085792236 | 0.025758486 | 0.35466832 | 0.37857985 |
| **NK_2** | 0.41701102 | 1.9936589 | 0.6910794 | 0.14645559 | 0.992215 | 0.6515524 | 0.79225105 | 0.1264289 | 1.9965264 | 2.1315467 |
| **NK_3** | 0.34367085 | 1.8145276 | 0.8420558 | 0.14902283 | 0.88689864 | 0.59097034 | 0.7207655 | 0.08296705 | 1.2739339 | 2.2210832 |
| **Naive B** | 0.002660668 | 0.07743975 | 0.001100029 | 0.002715168 | 0.01758262 | 0.004457866 | 0.009268549 | 0.000895799 | 0.41441116 | 0.035816487 |
| **Naive CD4 T** | 0.003025895 | 0.13266796 | 0.001389389 | 0.001210414 | 0.10300078 | 0.00732378 | 0.064799 | 0.028871514 | 0.33728424 | 0.05057629 |
| **Naive CD8 T** | 0.00572027 | 0.14709277 | 0.002817698 | 0.000790966 | 0.1583283 | 0.008758929 | 0.03941298 | 0.007832052 | 0.31856588 | 0.05113922 |
| **Neutrophil** | 0.001352365 | 0.05906428 | 0.104667924 | 0.029206187 | 0.039260313 | 0.008851456 | 0.020986885 | 0.001768525 | 0.034489326 | 0.098766774 |
| **Non-classical monocyte** | 7.54E-05 | 0.095180385 | 0.00191182 | 0.004418194 | 0.40522194 | 0.000885516 | 0.007579541 | 8.36E-05 | 0.046262342 | 0.12377018 |
| **Plasma cell** | 0.000295047 | 0.21627088 | 0.001977517 | 0.00095193 | 0.06390788 | 0.003337974 | 0.007771222 | 0.001286547 | 0.20007893 | 0.115437336 |
| **Plasmacytoid DC** | 0 | 0.11294102 | 6.23E-06 | 0.005538434 | 0.049926743 | 0.011940355 | 0.000170203 | 0.000340467 | 0.110636435 | 0.026059398 |
| **Platelet** | 0.000369337 | 0.12966199 | 0.002453423 | 0.004276724 | 0.04896173 | 0.037015304 | 0.005681763 | 0.001702778 | 0.22413704 | 0.08264785 |
| **Undefined** | 0.000201917 | 0.2150537 | 0.000257167 | 0.004077684 | 0.09967305 | 0.006564809 | 0.00217415 | 0.002712003 | 0.38672838 | 0.11175917 |
| **hNK1** | 0 | 1.6272125 | 0.002581815 | 0.000552984 | 0.6704472 | 0.001175255 | 0 | 0.000386713 | 0.47642884 | 0.5495319 |
| **hNK2** | 0 | 1.6115638 | 0.000280271 | 0 | 0.7951309 | 0.000378525 | 0 | 0.000135928 | 0.46123427 | 0.75215715 |
| **hNK3** | 0 | 1.4153008 | 0.000718241 | 0.000242132 | 0.49616864 | 0.001703944 | 2.46E-05 | 0.000420454 | 0.3834582 | 0.4917156 |
| **hNK4** | 0 | 1.446101 | 0.000143007 | 0 | 0.594497 | 0.001275949 | 0 | 0.000860135 | 0.5831781 | 0.6811563 |
| **hNK5** | 0 | 1.1157582 | 0 | 0 | 0.27004194 | 0.003661398 | 0 | 0.004735214 | 0.5942831 | 0.6704164 |
| **hNK6** | 0 | 1.1480199 | 0.000550072 | 0.000337627 | 0.3920326 | 0.002004543 | 4.71E-05 | 0.001398855 | 0.43254092 | 0.47037616 |
| **mNK1** | 1.5699917 | 4.3010306 | 1.8104784 | 1.0404555 | 2.1989927 | 1.8378602 | 1.8980232 | 0.5034612 | 2.9446058 | 4.182051 |
| **mNK2** | 1.2750257 | 3.6494646 | 1.6269764 | 0.49560258 | 1.8560386 | 1.5847834 | 1.496086 | 0.3654902 | 3.145605 | 4.757742 |
| **mNK3** | 0.5861804 | 2.5469663 | 0.9171339 | 0.1671475 | 1.190462 | 0.85160494 | 1.0150831 | 0.2720078 | 2.3901217 | 3.2926044 |
| **mNK4** | 0.6674098 | 2.7888274 | 1.0690004 | 0.17158559 | 1.3737108 | 1.0159653 | 1.1414486 | 0.28033566 | 2.713107 | 3.8877246 |
| **mNK5** | 0.6498435 | 2.8841152 | 1.0100619 | 0.25403455 | 1.2053277 | 0.9002229 | 1.225386 | 0.54986775 | 2.741384 | 4.0462933 |
| **mNK6** | 1.2408471 | 3.3191946 | 1.528023 | 0.56557333 | 1.6818576 | 1.5265431 | 1.417667 | 0.5765113 | 2.3820438 | 3.5895445 |
| **mNK7** | 0.6145842 | 2.1253743 | 0.4506978 | 0.15221752 | 0.3923917 | 0.4772011 | 0.84848076 | 0.55684376 | 2.3403373 | 2.0983093 |
