## Supplementary Table 3 for "Terminally differentiated CX3CR1^+^ natural killer cells drive coronary arteritis through an NFAT–T-bet axis"

**Supplementary table 3. Human scRNAseq QC**

| Sample | GEO sample (GSM) | Patient / donor ID | GEO series | Group | Max detected genes(nFeature_RNA <) | Max UMI counts(nCount_RNA <) | Max mitochondrial reads(percent_MT < , %) | Min cells per gene | Min genes per cell | Genes after QC | Cells after QC |
|---|---|---|---|---|---|---|---|---|---|---|---|
| KD1 | GSM5091060 | P53 | GSE167029 | KD | 4000 | 20000 | 15 | 3 | 200 | 14747 | 8900 |
| KD2 | GSM5160417 | P1 (before IVIG) | GSE168732 | KD | 2000 | 6000 | 5 | 3 | 200 | 15137 | 4762 |
| KD3 | GSM5160420 | P2 (before IVIG) | GSE168732 | KD | 4000 | 10000 | 5 | 3 | 200 | 16047 | 5213 |
| KD4 | GSM5160422 | P3 (before IVIG) | GSE168732 | KD | 4000 | 15000 | 5 | 3 | 200 | 16284 | 5455 |
| KD5 | GSM5160424 | P4 (before IVIG) | GSE168732 | KD | 4000 | 15000 | 5 | 3 | 200 | 16683 | 5197 |
| KD6 | GSM5160427 | P5 (before IVIG) | GSE168732 | KD | 4000 | 15000 | 5 | 3 | 200 | 16646 | 8567 |
| KD7 | GSM5160430 | P6 (before IVIG) | GSE168732 | KD | 4000 | 15000 | 5 | 3 | 200 | 15778 | 3848 |
| KD8 | GSM6042979 | KD1053 | GSE200743 / GSE254657 | KD | 4000 | 20000 | 7 | 3 | 200 | 17313 | 4845 |
| KD9 | GSM6042980 | KD1092 | GSE200743 / GSE254657 | KD | 5000 | 20000 | 7 | 3 | 200 | 17468 | 3706 |
| KD10 | GSM6470208 | KD1340 | GSE200743 | KD | 4000 | 10000 | 5 | 3 | 200 | 16487 | 5360 |
| KD11 | GSM8047972 | KD1278-1 | GSE254657 | KD with CAL | 5000 | 20000 | 7.5 | 3 | 200 | 20433 | 5919 |
| KD12 | GSM8047973 | KD1408-1 | GSE254657 | KD with CAL | 5000 | 15000 | 7.5 | 3 | 200 | 24009 | 15698 |
| KD13 | GSM8047974 | KD1409 | GSE254657 | KD with CAL | 5000 | 20000 | 7.5 | 3 | 200 | 23081 | 6290 |
| HC1 | GSM5073064 | C.HD1 | GSE166489 | Healthy | 3500 | 15000 | 5 | 3 | 200 | 17618 | 9238 |
| HC2 | GSM5073065 | C.HD2 | GSE166489 | Healthy | 2500 | 7500 | 7.5 | 3 | 200 | 16303 | 5856 |
| HC3 | GSM5073066 | C.HD3 | GSE166489 | Healthy | 3000 | 15000 | 5 | 3 | 200 | 17435 | 9607 |
| HC4 | GSM5073067 | C.HD4 | GSE166489 | Healthy | 3000 | 10000 | 5 | 3 | 200 | 17318 | 10693 |
| HC5 | GSM5073068 | C.HD5 | GSE166489 | Healthy | 3000 | 10000 | 5 | 3 | 200 | 16876 | 11260 |
| HC6 | GSM5073069 | C.HD6 | GSE166489 | Healthy | 3000 | 10000 | 5 | 3 | 200 | 16539 | 9324 |
| HC7 | GSM5160432 | H1 | GSE168732 | Healthy | 3000 | 10000 | 5 | 3 | 200 | 16128 | 5371 |
| HC8 | GSM5160434 | H2 | GSE168732 | Healthy | 3000 | 10000 | 5 | 3 | 200 | 15985 | 4682 |
| HC9 | GSM5160435 | H3 | GSE168732 | Healthy | 3000 | 10000 | 5 | 3 | 200 | 16041 | 5172 |
| HC10 | GSM8047977 | HC2790 | GSE254657 | Healthy | 5000 | 20000 | 7.5 | 3 | 200 | 25453 | 14190 |
| HC11 | GSM8047978 | HC2825 | GSE254657 | Healthy | 6000 | 20000 | 7.5 | 3 | 200 | 24749 | 12575 |
